# *De novo* Design of Macrocyclic Molecular Glues

**DOI:** 10.64898/2026.08.21.746227

**Authors:** Andrä Brunner, Krzysztof Wierbilowicz, Diandra Daumiller, Daniel Bexell, Kasper Karlsson, Olle Sangfelt, Patrick Bryant

## Abstract

The engineering of induced proximity has transformed drug discovery, yet the development of molecular glues remains largely serendipitous and restricted to the retrospective optimisation of accidental discoveries. Here, we present EvoBind-multimer, a deep learning framework for the *de novo* design of molecular glues directly from protein sequences. Unlike structure-based docking, our method generates small macrocyclic peptides that bridge user-defined protein pairs without requiring prior interface knowledge or existing ligands. We applied this framework to recruit the E3 ligase VHL to two challenging oncoproteins: KRAS and BRD4. Live-cell NanoBRET demonstrated robust design-induced proximity for both pairs. Mechanistic validation demonstrated that the generated macrocycles form functional VHL-target ternary complexes capable of driving Cullin-RING ligase-dependent proteasomal degradation and downstream signalling shutdown. Finally, evaluation in patient-derived xenograft neuroblastoma tumoroids revealed that ternary complex processing is deeply context-dependent: identical macrocycles acted as potent degraders in one patient model, yet functioned as stabilising “LOCKTACs” in another, driving VHL-dependent target sequestration without turnover. By enabling the *de novo* design of induced proximity from sequence alone, EvoBind-multimer provides a route towards designing new protein functions.

## Introduction

The engineering of novel protein-protein interactions (PPIs) represents a fundamental challenge in synthetic biology and therapeutics. While traditional drug discovery has excelled at designing molecules that inhibit protein function by blocking active sites, creating de novo interactions that induce proximity between two non-interacting proteins remains a formidable challenge. Mastering this capability would allow us to rewire cellular circuitry at will, opening vast possibilities for modulating biological systems beyond simple inhibition.

This paradigm of “pharmacological proximity” extends far beyond simple degradation. By rationally bridging any two proteins, we could direct specific enzymes to novel substrates, e.g. recruiting phosphatases to deactivate signalling pathways, or kinases to activate them. Furthermore, it enables precise spatial control, such as sequestering targets to specific organelles via Lysosome-Targeting Chimaeras (LYTACs) [1] or LOCTACs [2] for organelle-specific degradation. However, realising this vision requires a generalisable technology capable of designing interfaces between arbitrary proteins, a capability that currently does not exist.

Currently, the most transformative application of this proximity concept is Targeted Protein Degradation (TPD) [3]. Unlike inhibitors that require high systemic exposure to maintain receptor occupancy, TPD leverages the cell’s innate quality control machinery. By recruiting an E3 ubiquitin ligase to a specific target protein, these engineered interactions facilitate the formation of a ternary complex [4]. This induced proximity enables ubiquitination of the target, tagging it for irreversible proteasomal destruction [5], [6]. While TPD serves as the primary testbed for proximity-inducing therapeutics, the principles of ternary complex formation apply universally to other modalities, such as those targeting viral infections or autoimmune diseases [7], [8].

Despite this potential, the field is currently dominated by PROTACs (Proteolysis-Targeting Chimaeras), which face significant structural and design limitations. PROTACs are typically bifunctional molecules constructed by chemically linking a known effector binder (e.g., for an E3 ligase) to a known target binder. This “stitch-and-sew” approach relies heavily on the availability of pre-existing binders for both proteins, severely restricting the druggable proteome. Furthermore, the optimisation of bifunctionals is laborious [9] and subject to “hook effects” [10]. Consequently, while the human genome encodes over 600 E3 ligases [11], only a tiny fraction are currently exploitable due to a lack of known chemical handles.

A more elegant solution, applicable to both TPD and broader proximity applications, lies in molecular glues. These are monovalent molecules that stabilise the interface between two proteins de novo, without the need for linkers. Numerous molecular glues are currently employed in the clinic, most notably the immunomodulatory imide drugs (IMiDs) such as thalidomide and lenalidomide, which predominantly engage the E3 ligase cereblon (CRBN) [12]. However, virtually all clinically approved molecular glues were discovered entirely by serendipity or retrospective phenotypic screening rather than through rational design [13,14].

The rational design of molecular glues requires precise structural orchestration to bridge two proteins that have no natural affinity for one another. Current methods are hindered by their reliance on pre-existing high-resolution structures (holo structures). This strategy is insufficient because the formation of a ternary complex frequently induces significant conformational changes, remodelling the protein surfaces to create novel binding interfaces that do not exist in isolation. Therefore, effectively targeting these dynamic states requires a fully flexible, sequence-based framework capable of co-folding the components to capture induced-fit interactions, rather than merely docking onto rigid templates. Here, we present a computational framework for the de novo design of macrocyclic molecular glues directly from sequence information, validate their target degradation, and demonstrate efficacy in patient-derived xenograft tumoroids.

## Results

### The EvoBind-multimer Design Framework

We present EvoBind-multimer (EBM), a novel generative method for designing macrocyclic peptides acting as molecular glues (MGs) to facilitate interactions between two different proteins. EBM requires only the amino acid sequences of two target proteins and generates a cyclic peptide sequence-structure combination that binds to both simultaneously, thereby facilitating their interaction **(Figure 1).** EBM identifies suitable binding regions on each protein and orchestrates the interfaces between them without any prior structural information, constituting a blind, *de novo* design process.

**Figure 1.**
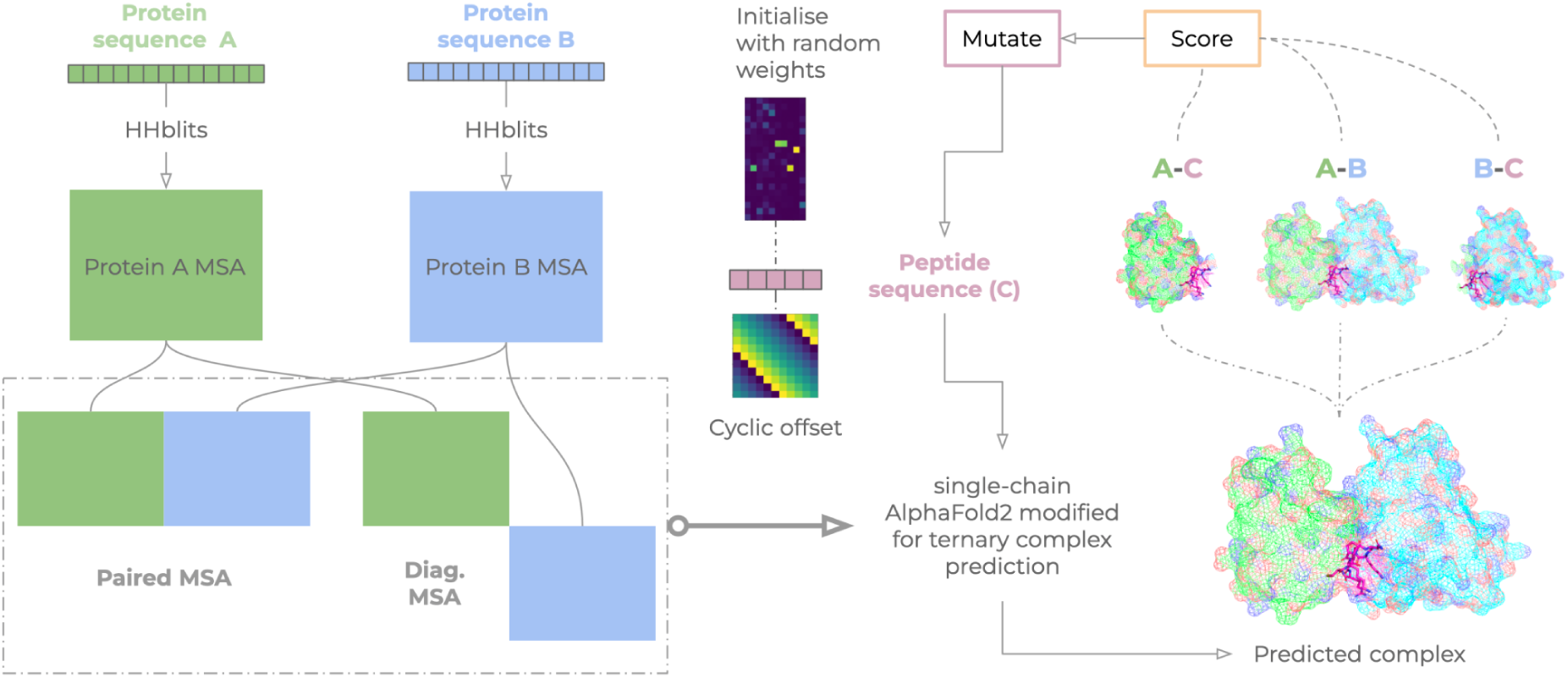
EvoBind-multimer (EBM) designs macrocyclic peptides to facilitate the interaction between two target proteins using only their sequences. The EBM workflow begins with the sequences of two distinct target proteins, Protein A (green) and Protein B (blue). Multiple Sequence Alignments (MSAs) are generated with HHblits for each protein and are subsequently processed to create paired and diagonalised MSA representations. These MSAs, which capture co-evolutionary information, are used as input for a single-chain AlphaFold2 model modified for ternary complex prediction. The peptide design process is an iterative optimisation loop. First, a macrocyclic peptide sequence (C, magenta) is initialised with random weights and a cyclic offset. This peptide, along with the target protein MSAs, is fed into the modified single-chain AlphaFold2 model to predict the structure of the full ternary complex (A-B-C). A scoring function then evaluates the quality of this predicted complex, assessing the pairwise interactions between the peptide and each target (A-C and B-C), and indirectly the baseline interaction between the two targets (A-B). The peptide sequence is then mutated, and the cycle of prediction and scoring is repeated. This evolutionary process optimises the peptide sequence to act as a “molecular glue,” resulting in a final, high-confidence predicted ternary complex where the designed peptide bridges Protein A and Protein B.

To achieve this, we modified single-chain AlphaFold2 (AF) to process data from three distinct entities: (1) target A, (2) target B, and (3) the peptide binder acting as a bridge between them. The different chains are represented via artificial chain breaks, achieved by setting residue offsets between each sequence so that each component is treated as a separate physical entity [15]. This modification allows AF to model the complex holistically, even though the original network was not trained on multimeric structures [16]. We also analysed the possibility of using AlphaFold3 [17] and Boltz-2 [18] but found that these complex-biased models consistently produce faulty interfaces. These interfaces mimic the training data from the PDB, inconsistent with the findings here, thereby reporting low predicted confidence (**Supplementary Figures 1&2**). EBM is fundamentally a sequence-based framework, yet it is guided by rigorous structural and coevolutionary considerations. For each target protein sequence, a multiple sequence alignment (MSA) is generated. These MSAs provide EBM with insights into the coevolutionary relationships within and between the two target proteins, which the network leverages to predict the folded state of the protein-protein-peptide ternary complex.

The generative process operates via an iterative optimisation loop. Following an initial prediction, the framework evaluates the predicted confidence of the peptide (pLDDT, predicted local distance difference test [19]) alongside the atomic proximity between the peptide and the two target proteins (*equation 1*, Methods). If these structural metrics improve, we accept the new sequence as an updated starting point and introduce a random mutation at a single amino acid position (**Figure 1**). This directed mutation procedure runs for 1000 iterations, systematically evolving the peptide sequence until it converges on a stable, high-confidence ternary complex.

### Macrocyclic Glue Design for VHL-KRAS and VHL-BRD4

To demonstrate the potential of EBM real-world applications, we applied the framework to the design of macrocyclic peptides that recruit the VHL E3 ubiquitin ligase [20] to two critical cancer targets: KRAS [21] and BRD4 [22]. By generating peptides that bridge these proteins, EBM effectively designs novel MGs [23] *de novo*, using only protein sequence information as input **(Figure 2a),** for the purpose of degrading the target proteins **(Figure 2b)**. To maximise potential for cell permeability, we restricted the design space to small macrocycles (4-10 residues). However, it is inherently more difficult to design such small macrocycles compared to larger proteins or bulkier scaffolds; limiting the sequence length drastically restricts the structural real estate available to establish a stable, high-affinity interfacial footprint capable of productively bridging two distinct protein surfaces.

**Figure 2.**
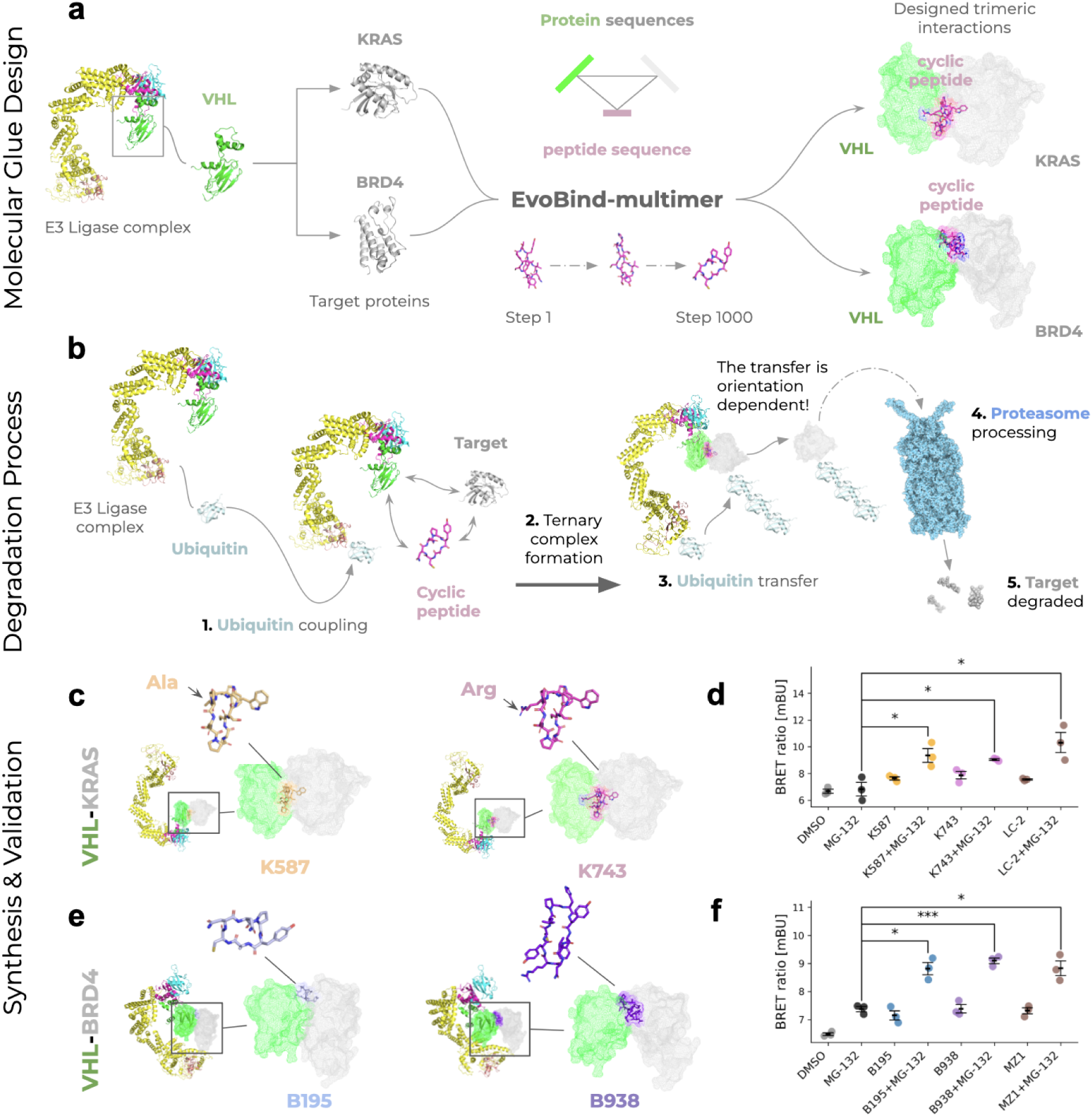
Design and functional validation of macrocyclic degraders targeting KRAS and BRD4. **a)** Schematic of the EvoBind-multimer (EBM) design framework applied to molecular glue (MG) design. EBM takes the amino acid sequences of an E3 ligase receptor (here, VHL; green) and a target protein (KRAS or BRD4; grey) as sole inputs. Using a structure-free, sequence-based evolutionary strategy (centre), it generates macrocyclic peptides (magenta) optimised to simultaneously bind both proteins and stabilise a ternary complex. **b)** Theoretical mechanism of targeted protein degradation induced by the designed macrocycles. **(1)** Ubiquitin is coupled to the E3 ligase. The peptide recruits the target to the E3 ligase complex, facilitating the formation of a productive ternary complex **(2)** and subsequent ubiquitin transfer **(3)**, which flags the target for proteasomal degradation **(4)**, resulting in the target protein being degraded **(5)**. **c)** Structural predictions of KRAS degraders K587 and K743. EBM predicted structures of the VHL-KRAS ternary complex mediated by the two top-ranked macrocyclic designs. The full E3 ligase complex (yellow/cyan), VHL (green), and KRAS (grey) are shown alongside the designed cyclic peptides K587 (orange, left) and K743 (magenta, right) (see **Table 1** for sequences and pLDDT confidence scores). **d)** NanoBRET quantification of KRAS ternary complex formation. Intracellular target engagement was assessed via NanoBRET assay. BRET ratios, expressed in milli-BRET units (mBU; calculated as 1000 × fluorescence signal/luminescence signal), are shown for cells treated with K587 and K743 compared to DMSO vehicle, the proteasome inhibitor MG-132 alone, and the PROTAC LC-2 positive control. Co-treatment with the proteasome inhibitor MG-132 results in significant accumulation of the complex. Data are represented as mean ± SEM (n=3 biological replicates). Statistical significance was determined using a two-sided unpaired Welch’s t-test (* p < 0.05). **e)** Structural predictions of BRD4 degraders B195 and B938. EBM structure predictions for VHL-BRD4 ternary complexes mediated by the designed macrocycles B195 (light blue, left) and B938 (purple, right) (sequences and pLDDT shown in **Table 1**). **f)** NanoBRET quantification of BRD4 ternary complex formation. Intracellular BRET ratios, expressed in milli-BRET units (mBU; calculated as 1000 × fluorescence signal/luminescence signal), demonstrating target engagement and complex formation for BRD4 degraders B195 and B938 compared to DMSO, the proteasome inhibitor MG-132, and the MZ1 PROTAC positive control. Measurements are shown both in the presence and absence of MG-132. Data are represented as mean ± SEM (n=3 biological replicates). Statistical significance was determined using a two-sided unpaired Welch’s t-test (* p < 0.05, *** p < 0.001).

**Table 1.** Selected EvoBind-multimer designs for VHL-KRAS and VHL-BRD4. The table lists top-ranking macrocyclic peptide candidates identified via the EvoBind-multimer pipeline. The peptides are named after their iteration number (e.g. K587 for KRAS, iteration 587). **Sequence** denotes the amino acid sequence of the designed macrocycle. **Length** indicates the number of residues. Computational design metrics include **pLDDT** (AlphaFold predicted Local Distance Difference Test confidence score), **loss** (final optimisation loss value, equation 1), and **IF distance** (interface distance metrics quantifying the proximity between the peptide and VHL or the target protein).

| VHL-KRAS |  |  |  |  |  |  |
| --- | --- | --- | --- | --- | --- | --- |
| iteration | sequence | length | IF distance VHL | IF distance KRAS | pLDDT | loss |
| 587 | APGDPVCSWW | 10 | 5,947 | 5,032 | 90,0 | 0,122 |
| 743 | RPGDPVCSWW | 10 | 5,797 | 5,547 | 90,5 | 0,125 |
| VHL-BRD4 |  |  |  |  |  |  |
| iteration | sequence | length | IF distance VHL | IF distance BRD4 | pLDDT | loss |
| 195 | YTCNNP | 6 | 4,915 | 4,940 | 91,3 | 0,108 |
| 938 | NRYCLPHYFV | 10 | 5,545 | 5,118 | 90,6 | 0,118 |

Overcoming this structural challenge, the EBM framework generated high-confidence small macrocycles for both target systems. To validate these designs experimentally, we synthesised two top-ranking candidates for each target system (comprising 6- and 10-residue peptides, **Table 1**) and evaluated their capacity to engage their targets and recruit the E3 ligase within cellular contexts. To circumvent permeability barriers during this proof-of-concept phase, peptides were administered via lipid-mediated intracellular delivery (Methods). We utilised a live-cell NanoBRET assay to monitor induced proximity through the energy transfer from NanoLuc luciferase linked to VHL to a fluorescent HaloTag conjugated to either KRAS or BRD4. Because highly active degraders can trigger rapid turnover of the ternary complex population, cells were evaluated in both the presence and absence of the proteasome inhibitor MG-132 [24] to allow the proximity signal to accumulate.

For the KRAS system, treatment with the 10-residue macrocycles K587 and K743 induced a clear increase in the BRET ratio, demonstrating successful intracellular ternary complex formation **(Figures 2c&d, Table 2)**. This proximity signal was significantly enhanced upon co-treatment with MG-132, which prevents proteasomal clearance of the ternary complex assembly, closely mirroring the trend observed for the positive control small-molecule PROTAC LC-2 [25]. Similarly, the BRD4-targeting macrocycles drove a robust and statistically significant accumulation of the BRET proximity signal under proteasome-inhibited conditions, tracking tightly with the established commercial benchmark PROTAC MZ1 [26] **(Figures 2e&f, Table 2)**. Collectively, these live-cell biophysical data demonstrate intracellular target engagement and VHL-dependent proximity induced by the *de novo*-designed macrocycles.

**Table 2.**
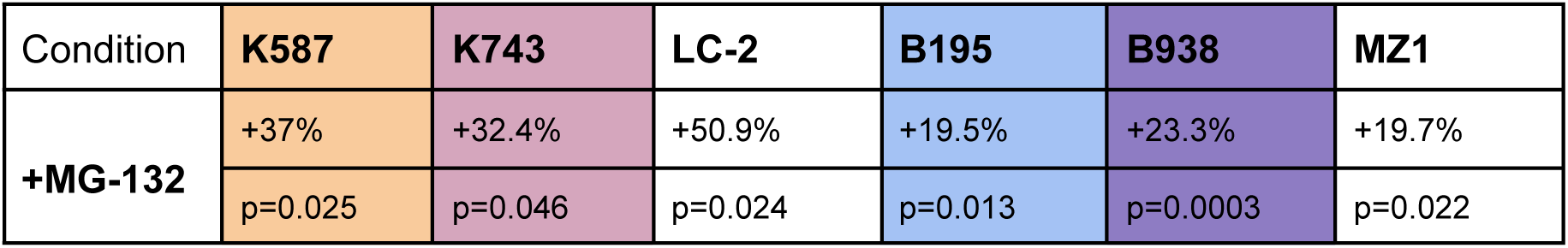
Live-cell NanoBRET ratios for VHL-KRAS and VHL-BRD4. The table lists top-ranking macrocyclic peptide candidates identified via the EvoBind-multimer pipeline. Target proximity was measured through excitation of the fluorescent Halo-tag on KRAS or BRD4 by luciferase linked to VHL. Cells were treated for 3 hours with 10 µM peptides, 1 µM LC-2 (KRAS +ctrl), or 100 nM MZ1 (BRD4 +ctrl), with or without a 1-hour pre-treatment of 10 µM proteasome inhibitor MG-132. Values represent the percentage change in BRET ratio (mBU) compared directly to the +MG-132 alone control baseline. Proteasome inhibition (+MG-132) results in strong ternary complex accumulation across all active compounds, suggesting that the macrocycles act as molecular glues. Statistical significance and the associated listed p-values were determined using a two-sided unpaired Welch’s t-test comparing the specified condition’s replicates directly to the replicates of the +MG-132 alone control group.

### Mechanistic validation of VHL and CRL-mediated target modulation

The recruitment of an E3 ligase is necessary but not sufficient for degradation; the ternary interface must enforce a specific geometry that aligns target surface lysines with the ubiquitin conjugate [27] (**Figure 2b**). If this geometric constraint is not met, the macrocycle may act as LOCKTAC rather than a degrader. To deconvolve the mechanism of target signal loss and distinguish between bona fide proteasomal degradation and non-functional steric sequestration, we interrogated the dependency of the macrocycle-induced phenotypes on the core ubiquitin-proteasome system (UPS) machinery. For the KRAS-targeting macrocycles (**Figure 3a**, **Table 3**), Western blot analysis confirmed a significant reduction in endogenous protein levels. Under baseline conditions, macrocycle 743 demonstrated robust degradation efficacy, clearing endogenous KRAS levels down to 0.3, which directly matched the performance of the established small-molecule PROTAC benchmark LC-2, while macrocycle 587 exhibited clearance to 0.5. This degradation effect was abrogated by siRNA-mediated knockdown of VHL (siVHL), which successfully rescued KRAS levels back to 0.7 for 743 and 0.6 for 587. Furthermore, inactivation of the Cullin-RING ligase (CRL) core via the neddylation inhibitor MLN4924 [28] successfully stabilised target levels for macrocycle 587 (0.5) and the LC-2 control (0.4). Together, these data confirm that macrocycle-driven KRAS clearance is explicitly driven by functional VHL recruitment and subsequent CRL2^VHL^-mediated proteasomal proteolysis.

**Figure 3.**
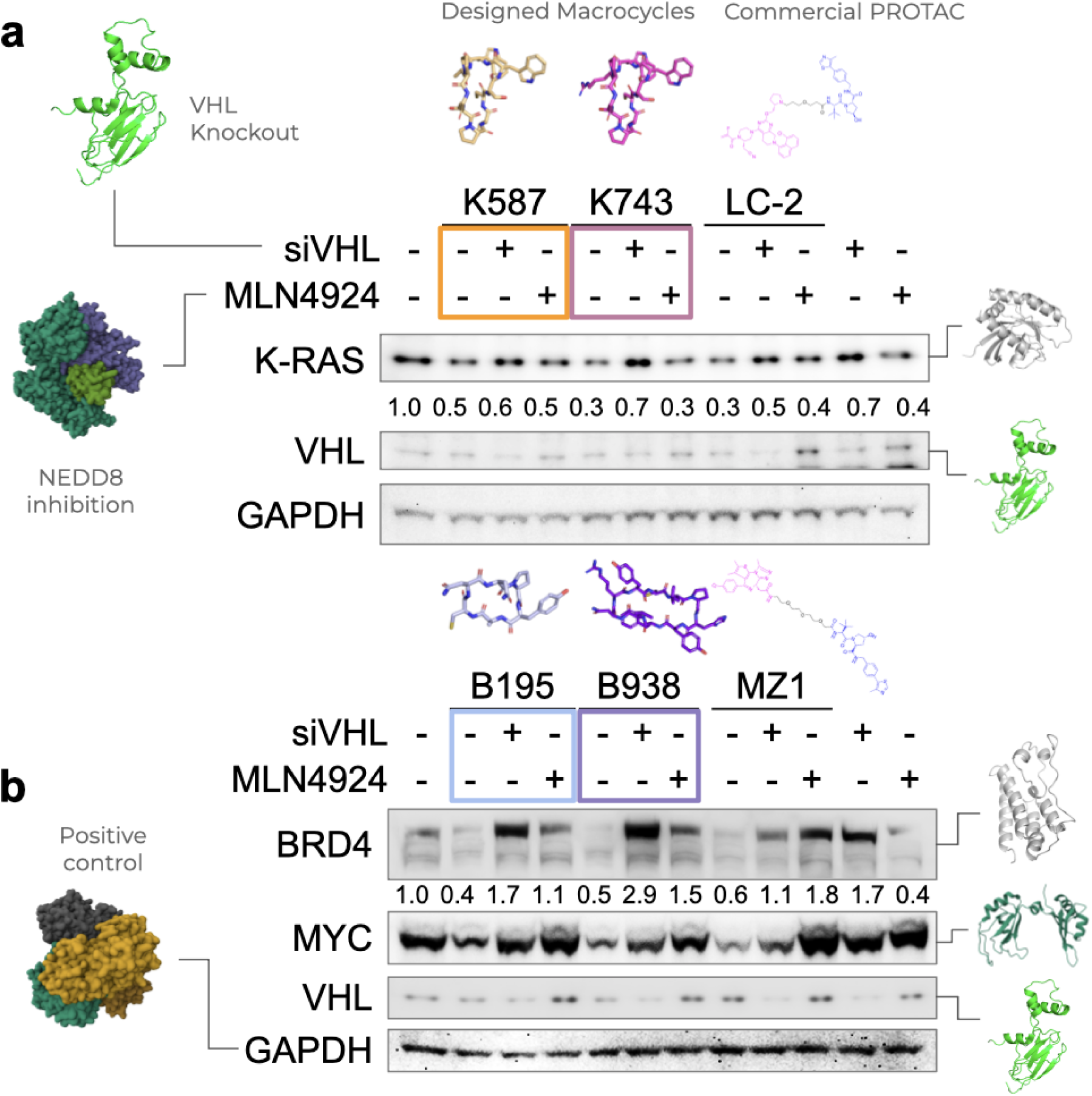
Mechanistic validation of VHL-mediated target modulation. Immunoblot analysis of endogenous target protein levels following 24 h treatment with vehicle control (DMSO), benchmark PROTAC controls (1 μM LC-2 or 100 nM MZ-1), or de novo-designed macrocycles (1 μM). Functional dependencies on the ubiquitin-proteasome pathway were mapped by profiling target abundance across modular pathway inhibition regimes. The matrix indicates pathway status: baseline conditions (−/−) represent vehicle; VHL depletion (+/−) indicates transfection with VHL-targeting siRNA 24 h before compound administration; and CRL inactivation (−/+) indicates pre-treatment with the NEDD8-activating enzyme (NAE) inhibitor MLN4924 (4 h before harvest) to block Cullin-RING ligase neddylation. GAPDH serves as a loading control. Representative VHL protein expression status validating siRNA-mediated knockdown efficiency and antibody reactivity is presented independently in the final two control lanes. Blots are representative of independent biological triplicates. Quantitation values below the target panels represent protein abundance normalised to the respective GAPDH loading control and relative to the DMSO baseline (1.0). **a) Mechanistic rescue of macrocycle-induced KRAS degradation.** Functional rescue assays performed under the unified pathway conditions outlined above utilising macrocycles 587 and 743 alongside the KRAS PROTAC benchmark LC-2. Under baseline conditions (−/−), macrocycle 587 exhibits moderate target degradation (abundance reduced to 0.5), while macrocycle 743 and the LC-2 benchmark demonstrate robust target clearance, both depleting endogenous KRAS levels down to 0.3. Depletion of VHL via siRNA (+/−) effectively compromises the degradation capacity of all three compounds, resulting in a marked rescue of KRAS protein levels to 0.6 for 587, 0.7 for 743, and 0.5 for LC-2. Inactivation of Cullin Neddyation via MLN4924 (−/+) yields target stabilisation values of 0.5 for 587 and 0.4 for LC-2, whereas the target abundance of macrocycle 743 remains at 0.3 within this specific kinetic window. See the Supplementary material for the full blots. **b) Mechanistic rescue of macrocycle-induced BRD4 degradation.** Parallel validation assays profiling BRD4 protein abundance and downstream MYC expression responsive to macrocycle treatment (B195 or B938) or the BRD4 PROTAC benchmark MZ-1. Under baseline conditions (−/−), macrocycles B195 and B938 demonstrate strong target degradation profiles, reducing BRD4 abundance to 0.4 and 0.5, respectively, while the MZ-1 benchmark clears the target down to 0.6. This clear reduction in BRD4 levels correlates with a corresponding downregulation of downstream oncogenic MYC expression across all three baseline treatment groups. Upon siRNA-mediated VHL silencing (+/−), macrocycle-induced degradation is abrogated, triggering a robust accumulation of BRD4 protein (1.7 for B195, 2.9 for B938, and 1.1 for MZ-1) and a concomitant stabilisation of downstream MYC protein levels. Under CRL inactivation via MLN4924 (−/+), BRD4 levels are similarly rescued to 1.1 for B195, 1.5 for B938, and 1.8 for MZ-1, confirming that macrocycle-driven target depletion operates via a highly cooperative, Cullin-RING-dependent proteasomal mechanism. See the Supplementary material for the full blots.

**Table 3.**
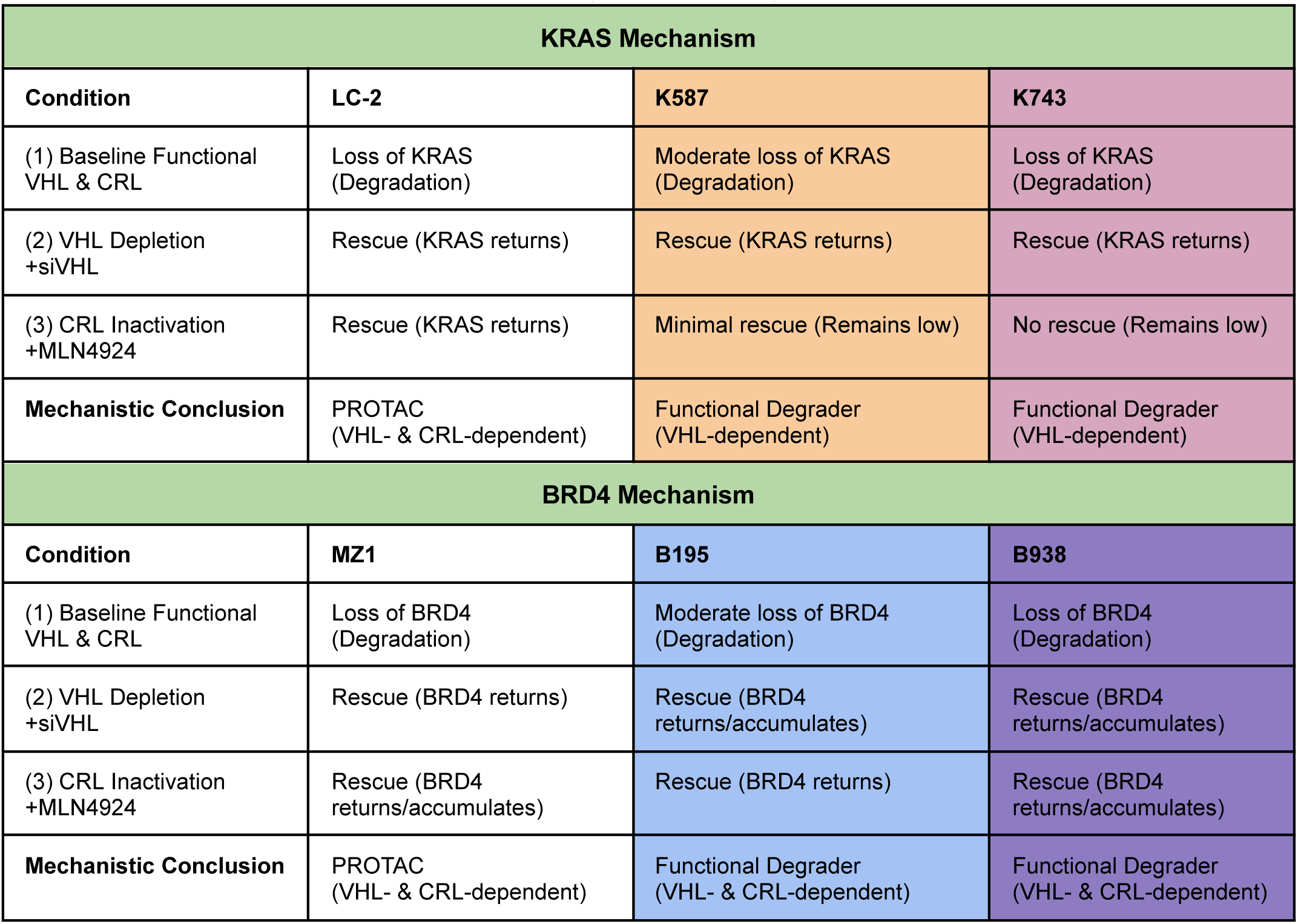
Mechanistic validation results and interpretation for the designed macrocyclic MGs. Summary of the mechanistic degradation profiles for the targeted KRAS and BRD4 degraders. Target protein levels were assessed under baseline conditions, following specific VHL knockdown (+siVHL) to test for ligase dependency, and upon global cullin-RING ligase (CRL) inactivation (+MLN4924). The results determine whether the observed target loss is driven by the anticipated VHL- and CRL-mediated proteasomal degradation pathways compared to established PROTAC controls (LC-2 and MZ1).

The mechanistic validation of the BRD4-targeting macrocycles (**Figure 3b**, **Table 3**) also demonstrated a genuine proteasomal degradation profile. Under baseline conditions, macrocycles B195 and B938 (1 μM each) drove intensive clearance of endogenous BRD4, reducing total protein abundance to 0.4 and 0.5, respectively, compared to 0.6 for the commercial benchmark MZ1 (100 nM). This target clearance robustly translated to a functional downstream biological shutdown, evidenced by a sharp downregulation of the oncogenic transcription factor MYC across both macrocycle treatment arms. To definitively map pathway dependencies, we evaluated target abundance in response to modular UPS blockade. siRNA-mediated depletion of VHL blocked macrocycle activity, triggering a dramatic hyper-accumulation of BRD4 (1.7-fold for B195 and 2.9-fold for B938 relative to the DMSO baseline) and a rescue of downstream MYC expression. Similarly, halting E3 ligase activation via MLN4924 rescued BRD4 levels to 1.1 and 1.5, respectively. Collectively, these Western blot rescue profiles across both target classes rule out non-functional sequestration, establishing our de novo-designed macrocycles as highly active, mechanically validated, Cullin-dependent degraders capable of driving robust target turnover and modulating downstream oncogenic signalling networks in human cells.

### Context-Dependent BRD4 Degradation and Sequestration in Patient-Derived Xenograft Neuroblastoma Tumoroids

To evaluate the therapeutic consequence of the designed BRD4-targeting macrocycles in physiological settings, we assessed their functional impact in patient-derived xenograft (PDX-derived) tumoroids of neuroblastoma, a cancer in which N-Myc expression is frequently regulated by BRD4 [29]. We hypothesised that VHL-mediated ubiquitination and subsequent degradation of BRD4 would abolish its chromatin-reading functions, effectively acting as a potent transcriptional repressor of oncogenic drivers. We evaluated two different PDX-derived models, LU-NB-2 and LU-NB-3 **(Figure 4a)** [30].

**Figure 4.**
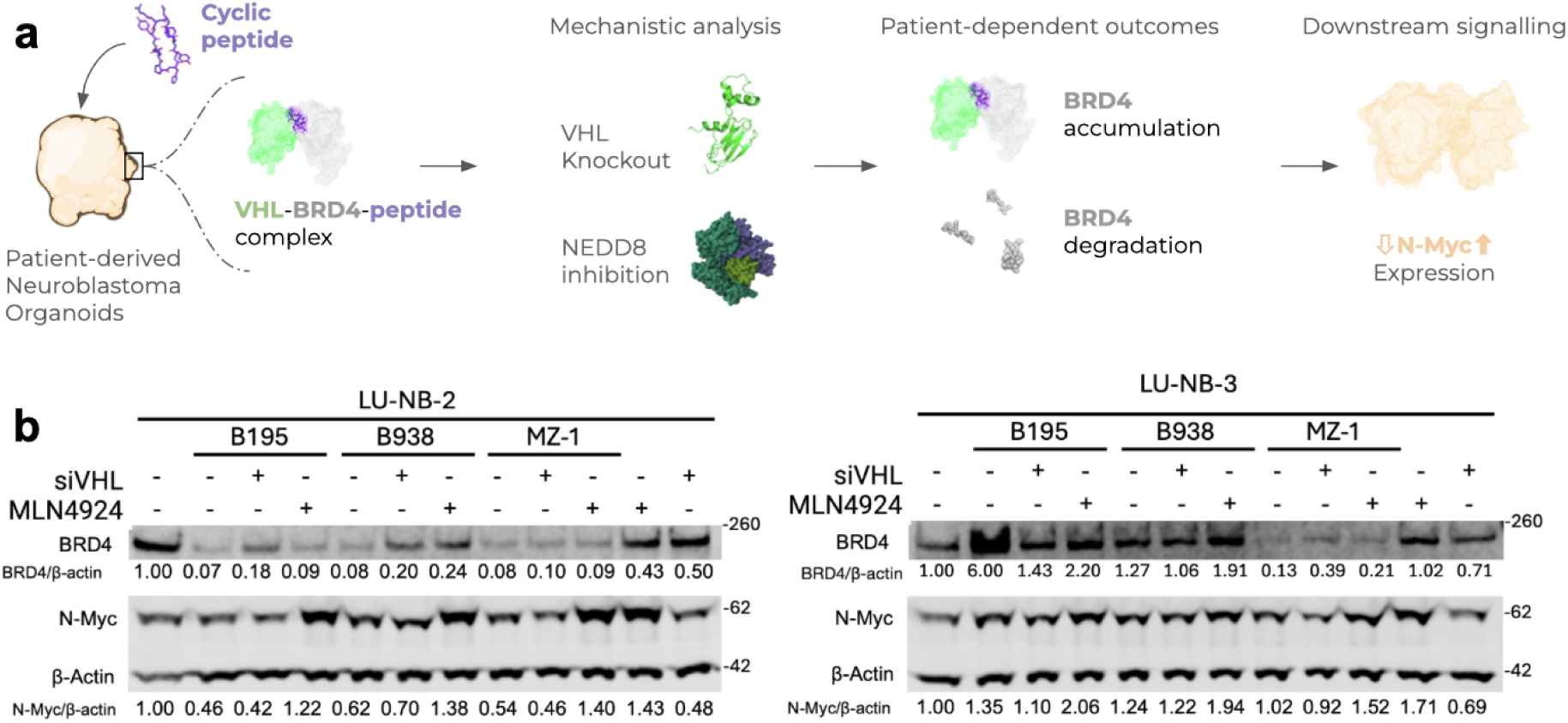
Patient-derived xenograft (PDX)-derived neuroblastoma tumoroids exhibit context-dependent degradation or stabilisation of BRD4. **a)** Schematic overview of the experimental workflow in 3D primary models. PDX-derived neuroblastoma tumoroids were treated with designed cyclic peptides to induce the assembly of the VHL-BRD4 ternary complex. Mechanistic dependencies were evaluated using VHL knockdown (siVHL) and Cullin-RING ligase/NEDD8 inhibition (MLN4924), followed by the assessment of downstream N-Myc signalling. This pipeline reveals divergent, patient-dependent cellular outcomes ranging from target degradation to target accumulation. **b)** Immunoblot analysis of LU-NB-2 and LU-NB-3 tumoroids treated with 10 µM of macrocycles (B195 or B938) or the benchmark PROTAC MZ-1 (100 nM). In the LU-NB-2 model, B195 and B938 function as robust, VHL- and CRL-dependent degraders that effectively clear BRD4 and repress N-Myc expression. In contrast, identically treated LU-NB-3 tumoroids exhibit a divergent LOCKTAC-like mechanism, where macrocycle treatment (particularly B195) drives a profound VHL-dependent accumulation of BRD4 without initiating target turnover. Densitometry values for BRD4 and N-Myc, normalised to the β-actin loading control, are indicated below the respective blots.

In the LU-NB-2 tumoroid model, treatment with the designed macrocycles B195 and B938 induced a robust decrease in endogenous BRD4 protein levels, effectively matching the performance of the commercial PROTAC benchmark MZ1 [26] **(Figure 4b)**. This degradation was directly dependent on the core ubiquitin-proteasome system machinery; target clearance was successfully rescued by siRNA-mediated depletion of VHL (siVHL) for B938 and pharmacological inactivation of the Cullin-RING ligase core via MLN4924 for both peptides. This targeted physical depletion of BRD4 was accompanied by a concurrent reduction in downstream N-Myc expression, with N-Myc/β-actin ratios dropping to 0.46 and 0.62 relative to baseline following treatment with B195 and B938, respectively. This repression closely mirrored the effect achieved by the MZ1 control (0.54). These data demonstrate that the generatively designed glues successfully hijack the cellular degradation machinery to eliminate their target and modulate oncogenic signalling within complex, PDX-derived 3D architectures.

Interestingly, evaluating the macrocycles in a second, parallel-processed PDX-derived model, LU-NB-3, revealed a starkly divergent mechanistic response. Rather than inducing target clearance, the macrocycles—particularly B195—drove a robust accumulation of BRD4 protein. Importantly, this stabilisation effect was abrogated by siVHL. This confirms that the macrocycles successfully penetrate the cells and assemble the VHL-BRD4 ternary complex, but fail to trigger subsequent ubiquitination or proteasomal processing in this specific cellular environment, thereby converting the degrader into a VHL-dependent steric stabiliser (a LOCKTAC mechanism [2]). This profound phenotypic divergence between two identically handled primary patient models highlights a critical layer of cellular context-dependency in targeted protein degradation. Together, these data establish that EBM yields precision tools capable of engaging challenging targets in physiological tissues, while exposing complex, model-specific bottlenecks in ternary complex processing.

## Discussion

### From Serendipity to Programmable Macrocyclic Glue Design

Molecular glues modulate protein function through induced proximity, but serendipitous observations and retrospective optimisation have historically dominated their discovery. Existing rational approaches often depend on pre-existing binders, structural information or chemical starting points. Here, EvoBind-multimer (EBM) instead generates molecular glues directly from the amino acid sequences of two proteins, simultaneously designing the peptide and both protein-peptide interfaces *de novo*.

Rather than assembling two independent binders, EBM optimises the macrocycle within the complete ternary complex, directly exploring the space of three-way interfaces. This is particularly challenging for the small cyclic peptides generated here. Successful VHL-KRAS and VHL-BRD4 designs were only six to ten residues long, requiring EBM to simultaneously create two protein-peptide interfaces and their ternary arrangement within an exceptionally small molecular scaffold. Despite this severe size constraint, the resulting macrocycles were sufficient to induce intracellular proximity and target modulation.

EBM achieves this by repurposing the single-chain AlphaFold2 model to simultaneously model two target proteins and a cyclic peptide bridge. The model was not trained explicitly on the protein complexes, providing a route to interface design without requiring known protein-protein interactions. Consistent with this, AlphaFold3 and Boltz-2 preferentially generated interfaces resembling the canonical binding pockets common to known molecular glues and did not provide suitable predictions for the *de novo* interfaces explored here (Methods).

The successful experimental validation of EBM designs demonstrates that single-chain structure prediction can be repurposed to enable the *de novo* design of ternary interfaces mediated by very small macrocycles. More broadly, this suggests that models without an explicit prior for known protein interfaces may be advantageous when the objective is not to reproduce existing interactions, but to create new ones.

### Ternary complex geometry determines functional outcome

Our results highlight a critical feature of molecular glue design: E3 ligase recruitment is necessary but not sufficient for productive degradation. The ternary complex must adopt a geometry that permits efficient ubiquitination of the target. For KRAS, K587 and K743 induced intracellular VHL-KRAS proximity and reduced endogenous KRAS abundance, with degradation dependent on VHL and the CRL machinery, demonstrating that EBM can generate ternary complexes compatible with productive ubiquitination and degradation.

The BRD4 results further illustrate the importance of ternary complex processing. B195 and B938 produced VHL- and CRL-dependent BRD4 degradation in cells, accompanied by suppression of MYC. However, in the LU-NB-3 PDX-derived neuroblastoma tumoroids, the same macrocycles instead caused pronounced, VHL-dependent accumulation of BRD4. This contrasts with the degradation observed in the parallel LU-NB-2 model, where both macrocycles reduced BRD4 and N-Myc. Thus, the designed molecules can establish the VHL-BRD4 proximity in both cellular contexts, while the downstream fate of the ternary complex is context-dependent.

This distinction between programmable proximity and productive degradation is an important consideration for generative molecular glue design. Rather than optimising solely for ternary complex formation, future models could incorporate geometric objectives that favour specific downstream outcomes, such as positioning target lysines for ubiquitination [31]. More broadly, the LOCKTAC phenotype demonstrates that designed proximity can produce functional target modulation even when degradation does not occur, expanding the potential of EBM beyond targeted protein degradation.

### Outlook

Several challenges remain before EBM can be considered a general platform for molecular glue discovery. The present study establishes intracellular target engagement using lipid-mediated peptide delivery, and future work will need to address delivery and pharmacological properties independently of the design problem. More broadly, the success of the current designs does not imply that arbitrary protein pairs can yet be bridged efficiently. Expanding the range of targets, E3 ligases and cellular contexts will be important for establishing the generality of the approach.

Recent work has also demonstrated that large designed proteins can be used to bridge protein interfaces [32], highlighting the broader potential of generative approaches for proximity engineering. EBM extends this concept to a substantially smaller molecular format, generating macrocyclic peptides directly from the sequences of two target proteins without requiring pre-existing ligands, known binary interactions or experimentally determined ternary structures. The resulting molecules can form intracellular ternary interactions and induce target degradation, while the tumoroid experiments demonstrate that the same designed molecules can produce different functional outcomes depending on cellular context.

The broader implication is therefore not simply that molecular glues can be designed *de novo*, but that protein function may increasingly be engineered through the geometry of induced proximity. EBM provides a first step towards making ternary complex formation a computationally addressable design problem. Future generations of generative models may move beyond designing proximity itself towards designing the cellular outcome that proximity produces. After all, the ultimate goal is not proximity, but function.

## Methods

### Data

EvoBind-multimer (EBM) is an adaptation of the single-chain protein structure prediction method AlphaFold2 [16] and has therefore not seen any protein complex data. The generated complexes are thereby not based on explicit training examples of protein complexes, providing a test of the network’s capacity to generalise to a new design problem.

To test EBM, we selected the therapeutically important problem of targeted degradation [33] by bridging VHL-KRAS and VHL-BRD4. The computational complexity of EBM grows quadratically with the combined sequence length of the two target proteins. To reduce the computational demands, we selected ligase domains that are substrate-binding. In addition, we selected structural subsections of the ligase domains that do not have interfaces towards other subunits of the ligase. This is to guide the design procedure towards target interfaces that are not already occupied. **Table 4** provides an overview of the studied ligases, target proteins and selected regions. The full sequences and their lengths can be found in **Supplementary Table 1.**

**Table 4.** Overview of target systems VHL-KRAS and VHL-BRD4. In both cases, we use the wild-type sequences for design without introducing mutations.

| Ligase | Ligase domain | Target protein | Sequence lengths | PDB IDs |
| --- | --- | --- | --- | --- |
| von Hippel-Lindau (VHL)<br>Cullin RING E3 ligase | VHL:1-87 | KRAS: all residues | VHL: 139<br>KRAS: 170 | 8QU8 |
| von Hippel-Lindau (VHL)<br>Cullin RING E3 ligase | VHL:1-87 | BRD4: 5-134 (His5 in<br>beginning) | VHL: 139<br>BRD4: 134 | 8QU8 &<br>5Z1R |

### MSA and Peptide representations

For each sequence of the two target proteins to be bridged through binder design with EvoBind-multimer, we generated multiple sequence alignments (MSAs) using HHblits [34] version 3.1.0 and searching uniclust30_2018_08 [35] with the following options:

hhblits -E 0.001 -all -oa3m -n 2

From the resulting single-chain MSAs, we create paired and block-diagonalised versions [15] to extract potential coevolutionary relationships between the two target proteins. The pairing was done by taking the top hit for each UniProt OX (organism) identifier, and the block diagonalisation by adding gaps for the other chain(s) (**Figure 1).** The MSAs from the two target proteins are sampled together to obtain both coevolutionary signals from every single chain and between the pair (**Figure 1)**. Statistics for the different MSAs can be found in **Supplementary Table 2.**

The designed peptide binders are represented with a single sequence, which is initialised randomly with weights in the range 0-1 from a Gumbel distribution, since these do not have coevolutionary relationships with natural sequences, resulting in that MSAs can’t be generated. A cyclic offset is implemented to connect the peptide amino acids in a continuous cycle [36,37].

### Design procedure and Scoring

A structural prediction is generated from the MSA and peptide representations, and the resulting architecture, together with predicted confidence metrics, is used to score the suitability of the peptide sequence in bridging an interaction between the two target proteins. To formally evaluate the design, we calculate a custom loss according to *equation 1*. Based on the resulting score, accepted sequences serve as novel starting points and undergo a random mutation at a single amino acid position (**Figure 1**).

This mutation procedure is performed for 1000 iterations. To generate a diverse set of solutions, we ran five different initialisations for peptide lengths ranging from 4 to 10 amino acids, yielding a comprehensive candidate library (n = 35,000 per target). We intentionally restricted the peptides to a small size to increase their likelihood of traversing cell membranes, a vital property for intracellular applications such as targeted protein degradation.

Computationally, the framework is highly efficient. A single iteration takes 30 to 40 seconds on average, meaning a complete 1000-iteration design trajectory finishes in 8 to 11 hours on a single NVIDIA A100 GPU equipped with 40 GB of vRAM. By executing all design runs in parallel, the entire generative pipeline for a given target pair was completed overnight. Furthermore, these generative settings are highly modular and can be easily adapted to explore different peptide lengths, iteration depths, and initialisation parameters depending on specific design requirements.

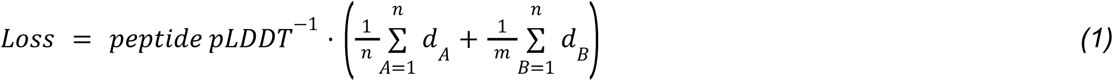

Where the peptide pLDDT is the average predicted lDDT [19] over the peptide, a measure of the local confidence of predicted protein structures (higher=better). The d_A_ and d_B_ are the shortest distances between all atoms n in the peptide and any atom in the two target proteins A and B.

### Design Loss and pLDDT Distributions

To evaluate the generative performance of the EBM framework, we analysed the distributions of the design loss and the predicted local distance difference test (pLDDT) scores across all generated macrocycles. For both the VHL-KRAS and VHL-BRD4 systems, we observed that longer peptide lengths were generally preferred by the network, consistently yielding lower loss values (**Figure 5**). This trend likely reflects the physical necessity of a larger interfacial footprint to effectively bridge two distinct, non-interacting protein surfaces. We note that larger structures are easier to design in general since information from larger structural regions can be used to generate high-confidence solutions. In contrast, a small peptide has a relatively small structural and coevolutionary impact on a ternary complex.

**Figure 5.**
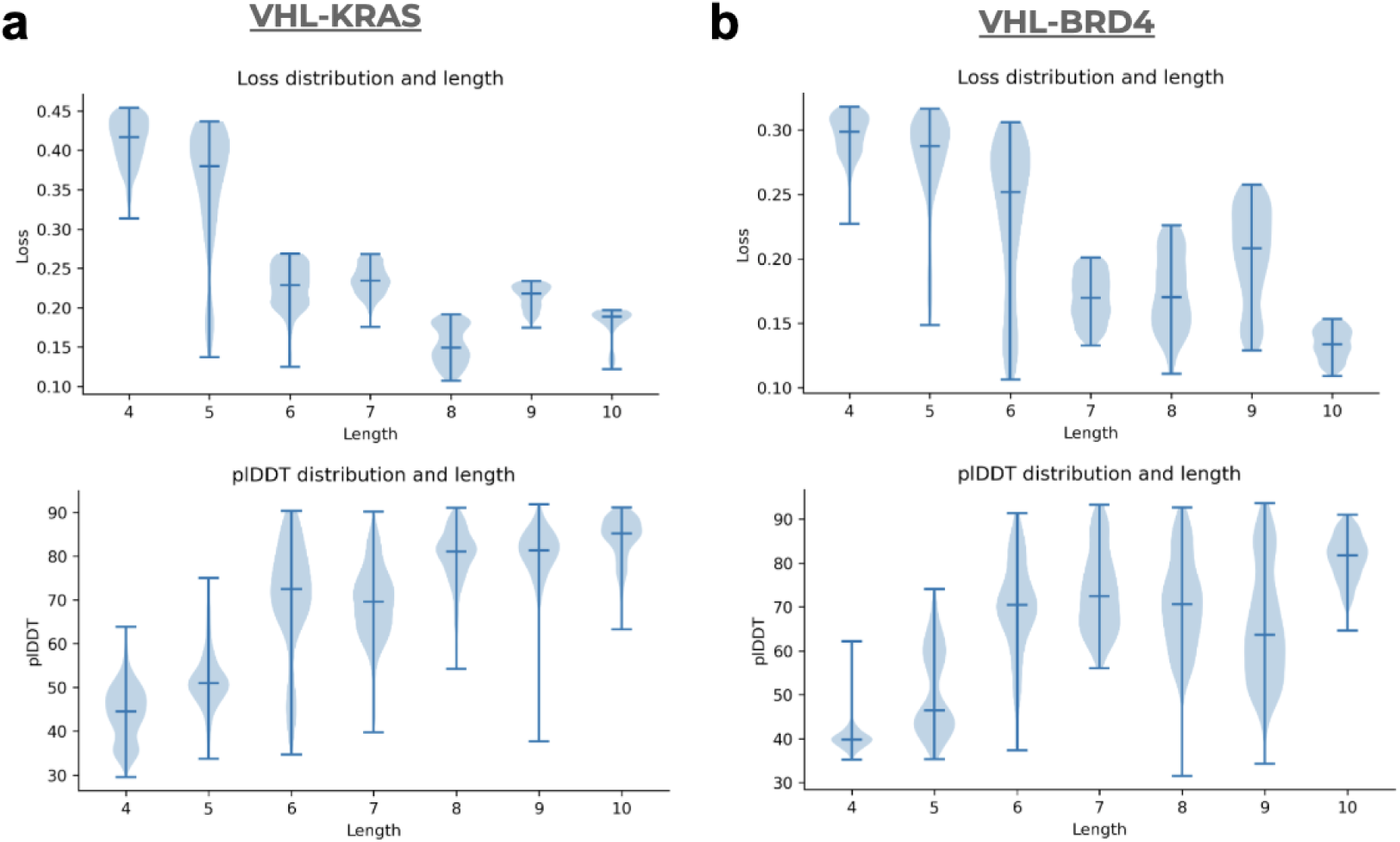
Computational evaluation of generative design metrics. Whiskers represent 95% confidence intervals, horizontal lines mark the medians. **a)** Distribution of design loss and peptide predicted local distance difference test (pLDDT) scores for macrocycles designed to bridge the VHL and KRAS target pair. The data represent the top 10% of generated sequences (n = 500 per peptide length) filtered according to *equation 1*. The generative framework demonstrates a clear physical preference for longer macrocyclic lengths, which correspondingly result in lower overall design losses and higher structural confidence. **b)** Corresponding design loss and peptide pLDDT distributions for macrocycles designed against the VHL and BRD4 target pair. Consistent with the KRAS dataset, the top 10% of generated sequences (n = 500 per length) exhibit improved generative metrics at larger peptide lengths, reflecting the increased interfacial surface area required to form a stable predicted ternary complex.

### Selection of Designs for Experimental Validation

We selected the top designs from all runs for the VHL target systems (VHL-KRAS and VHL-BRD4) according to a stepwise hierarchical filtering procedure.

First, we extracted the top 10% of generated sequences based on *equation 1* across all independent runs and peptide lengths, yielding an initial pool of 3500 candidate macrocycles per target system. Next, to ensure high structural fidelity of the predicted ternary complexes, we applied a strict confidence threshold, retaining only designs with a peptide pLDDT score ≥ 85. This reduced the candidate pools to 646 designs for VHL-KRAS and 353 designs for VHL-BRD4.

These pools were further narrowed down to the top 100 designs per target. An analysis of these subsets confirmed the network’s preference for larger macrocycles. For VHL-KRAS, the top 100 comprised 70 octapeptides (length 8) and 30 decapeptides (length 10). For VHL-BRD4, the distribution included 14 hexapeptides (length 6), 10 octapeptides (length 8), and 76 decapeptides (length 10).

Finally, we conducted a thorough analysis of the predicted structures of the remaining candidates. Since we focus on very short peptides, we did not perform an explicit selection based on aqueous solubility, given that most small cyclic peptides are inherently water soluble and computational estimations of this feature remain unreliable. For VHL-KRAS, we selected two peptides representing the best-performing length (10 residues). For VHL-BRD4, we selected two peptides corresponding to lengths of 6 and 10 residues to broadly sample diverse structural footprints. The final experimental candidates were explicitly chosen based on possessing the highest pLDDT scores, the lowest losses (equation 1), and absence of steric clashes upon manual visual inspection (**Table 1**).

### Ternary complex prediction with other AI models

AlphaFold-multimer (AFM) [38], AlphaFold3 (AF3) [17] and Boltz-2 [18] were used in a forward pass to re-predict the top four selected ternary complexes. This was done to both assess these models’ ability to model ternary complexes, as well as to potentially cross-validate the best designs. All predictions were run on a single NVIDIA A100 GPU equipped with 40 GB of RAM. Predicted ternary structures (AFM, AF3 and Boltz-2) were evaluated against the reference EvoBind-multimer models through Biopython [39]. The different models were superimposed onto the reference using Cα backbone atoms of the two protein target subunits (VHL and KRAS/BRD4) to calculate each protein target’s Cα RMSD (**Figure 6a**). Peptide RMSD was calculated across all non-hydrogen atoms of the cyclic peptide following the same structural alignment to account for both macrocyclic deformation and spatial displacement within the binding pocket (**Figure 6b**). The pLDDT comparison of these predictions are included in **Figure 6c and d**, while their structures are shown in **Supplementary Figures 1 and 2**. AF3 and Boltz-2 tend to reproduce interfaces found in the PDB, resulting in predictions very different from those observed with EBM. In addition, the AF3 and Boltz-2 predictions have low predicted confidence (pLDDT), possibly because the *de novo* designs don’t fit the interfaces the models have been exposed to during training. In the case of AFM, the predictions agree well with EBM for K587, showing high pLDDT. When the predictions do not agree as well (K743) or at all (BRD4) the pLDDT is much lower. This suggests that the models *“know”* that the interfaces they produce are not accurate, but they can’t find the ones generated by EBM.

**Figure 6.**
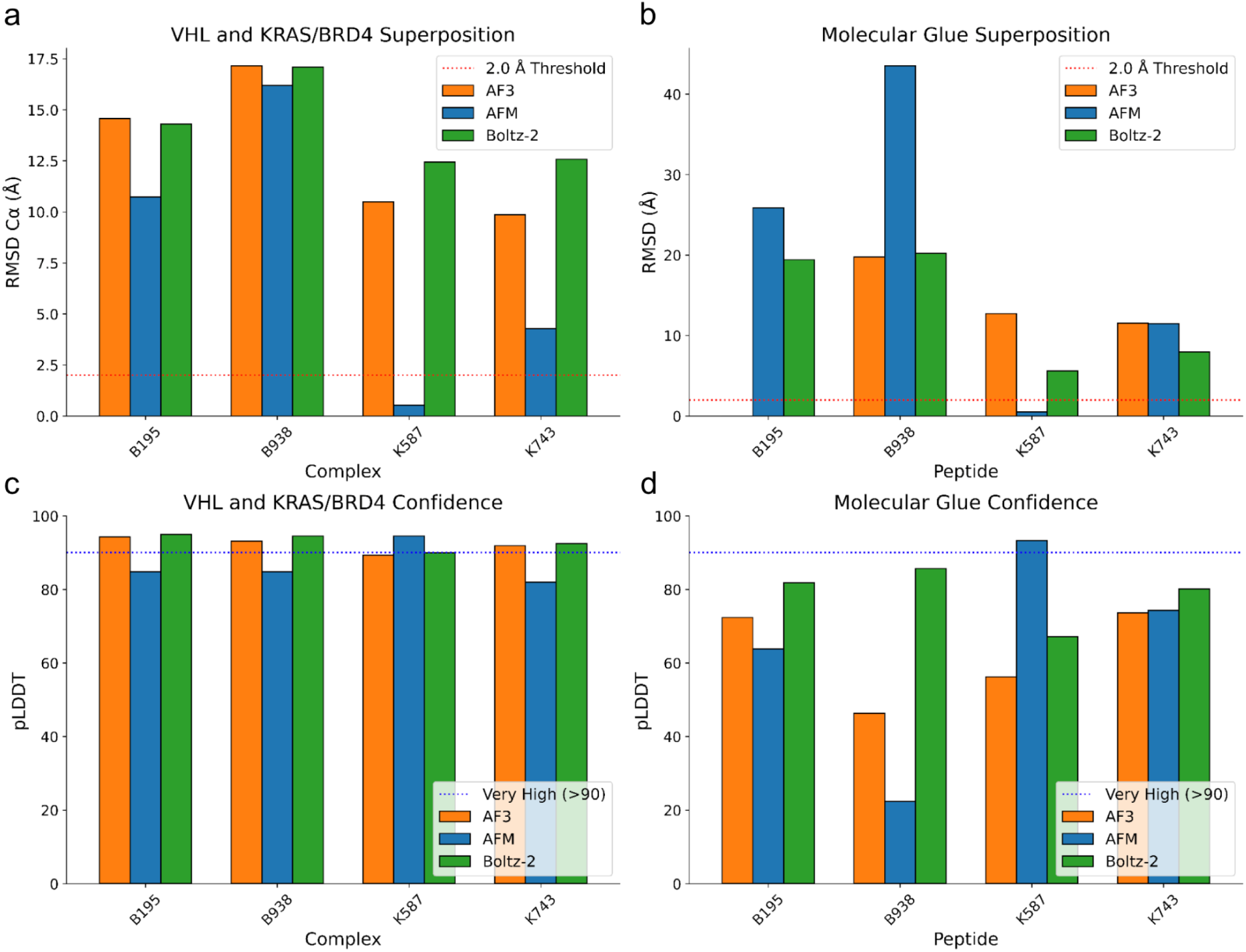
Structural accuracy and prediction confidence of the ternary complexes. Comparative evaluation of AF3 (orange), AFM (blue), and Boltz-2 (green) against the EvoBind-multimer reference model across the top four selected ternary complexes. **a)** Comparison of the protein target subunits VHL and KRAS or BRD4 Cα backbone atoms RMSD relative to the EBM reference. The dotted red line denotes the 2 Å threshold. **b)** RMSD comparison of the spatial displacement of all non-hydrogen heavy atoms of the cyclic peptide molecular glue against the reference EBM prediction. The dotted red line denotes the 2 Å threshold. **c)** Mean pLDDT score across the two protein target subunits Cα backbone atoms for the three cross-validation models. The dashed blue line represents highly confident (pLDDT > 90) predictions. **d)** Mean pLDDT score calculated across non-hydrogen heavy atoms of the cyclic peptide for the three cross-validation models. The dashed blue line represents highly confident (pLDDT > 90) predictions.

#### AFM ternary complex predictions

VHL and the target fasta sequences (KRAS and BRD4) were used as input to generate the AFM features required as input for AFM predictions. The peptide molecular glue sequences were input without MSA. For the prediction of ternary complexes, including cyclic peptides, AFM v2.1.0 was adapted and run with 20 recycles and dropout, but in the structural model (similarly to EvoBind2 [40]).

#### AF3 ternary complex predictions

Based on the same fastas and MSAs generated as input for EBM, a standard AF3 input JSON file (modelSeeds: [1–10], dialect: alphafold3, version: 2) was created, including the two protein targets as sequences and unpaired/paired MSAs (based on OX identifiers). Because standard protein sequence input pipelines in AF3 assume linear backbones and specifying covalent bonds within polymers is not supported (https://github.com/google-deepmind/alphafold3/blob/main/docs/input.md), the cyclic peptide glues were input as discrete ligand entities of Simplified Molecular-Input Line-Entry System (SMILES). SMILES for head-to-tail cyclic peptides were generated using RDKit (v2024.03.5 [41]) by constructing Hierarchical Editing Language for Macromolecules (HELM) strings with explicit N-to-C backbone cyclisation. HELM representations were parsed into molecular graphs via MolFromHELM and exported to canonical SMILES using MolToSmiles. AF3 was run using 20 recycles, and the top-ranked sample was used for RMSD comparisons.

#### Boltz-2 ternary complex predictions

The protein target sequences and unpaired HHblits MSAs were used in the Boltz-2 standard yaml input file. The cyclic peptide glue could also be directly specified as a cyclic protein with an empty MSA. Boltz-2 was run with 20 recycling steps and 2 diffusion samples per ternary complex. The first model (model 0) was then used for RMSD comparisons.

### Peptide purity and handling

Peptides were dissolved in the solvents according to **Table X** to create aliquots that were then used for all subsequent experiments. Both the purity and net peptide content were taken into account when calculating the molarity of the active peptide components. The final DMSO concentration in the cell assays was kept consistently at a maximum of 0.1% to avoid adverse effects and cell death.

**Table X.**
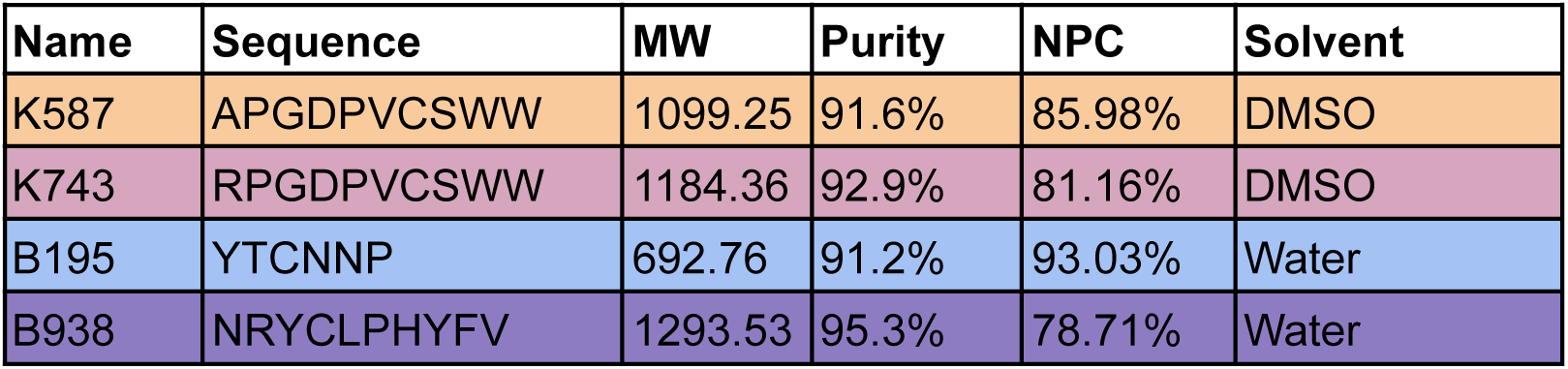
Molecular weight (MW), Purity, Net peptide content (NPC) and solvent used for each synthesised peptide.

| Name | Sequence | MW | Purity | NPC | Solvent |
| --- | --- | --- | --- | --- | --- |
| K587 | APGDPVCSWW | 1099.25 | 91.6% | 85.98% | DMSO |
| K743 | RPGDPVCSWW | 1184.36 | 92.9% | 81.16% | DMSO |
| B195 | YTCNNP | 692.76 | 91.2% | 93.03% | Water |
| B938 | NRYCLPHYFV | 1293.53 | 95.3% | 78.71% | Water |

### Cell culture and transfections

A549 cells were used for all KRAS-related assays and were transfected with peptides designed to bind KRAS and VHL. HCT116 cells were used for all BRD4-related assays and were transfected with peptides targeting BRD4 and VHL. Unless otherwise indicated, transfection conditions were identical across experiments and independent of the downstream readout, including western blot analysis, protein degradation reporter assays, immunoprecipitation, and cell viability assays. A549 and HCT116 cells were maintained in Dulbecco’s modified Eagle’s medium (DMEM) supplemented with 10% FBS and 2 mM L-glutamine, and incubated at 37 °C in 5% CO₂.

For peptide delivery, cells were transfected using the SAINT-Protein (Synvolux) transfection reagent according to the manufacturer’s instructions. For gene silencing, cells were transfected with a DsiRNA duplex (Integrated DNA Technologies) targeting VHL (sense: 5′-GAUUUCUGUUGAAACUUACACUGTT-3′; antisense: 5′-UUGUCACAUUCAAAGUUGUCUUUAG-3′) or a non-targeting control using HiPerFect (Qiagen) according to the manufacturer’s instructions. Cells were transfected with peptides 24 h before analysis or functional assays, and with siRNAs 48 h before analysis or functional assays.

### Live-cell NanoBRET assay

Proximity between VHL and either KRAS (G12V) or BRD4 was assessed using a live-cell NanoBRET assay. VHL was expressed as a C-terminal NanoLuc fusion (VHL-NanoLuc), whereas KRAS (G12V) and BRD4 were expressed as N-terminal HaloTag fusions (HaloTag-KRAS (G12V) and HaloTag-BRD4, respectively).

HEK293T cells were co-transfected with plasmids encoding VHL-NanoLuc and either HaloTag-KRAS (G12V) or HaloTag-BRD4 at a donor-to-acceptor plasmid ratio of 1:50 (VHL-NanoLuc:HaloTag fusion), using the transfection procedure described above. At 24 h after transfection, cells were replated in white 96-well plates at 5000 cells per well. NanoBRET HaloTag 618 ligand (Promega) was added to all experimental wells. A separate row of wells prepared without HaloTag 618 ligand was included for each VHL-target pair to determine the background emission ratio.

At 20 h after replating, macrocyclic peptides were introduced into cells by lipid-mediated delivery as described above. Cells were simultaneously treated with either 10 µM MG-132 (Selleckchem, S2619) or the corresponding vehicle. LC-2 (Selleckchem, S6996) and MZ1 (Selleckchem, S8889) were included as positive controls for the KRAS and BRD4 systems, respectively. After incubation for 4 h at 37°C, NanoBRET Nano-Glo substrate (Promega) was added according to the manufacturer’s instructions. Donor emission at 460 nm and acceptor emission at 618 nm were measured using a Spark multimode microplate reader (Tecan).

The raw BRET ratio was calculated by dividing the acceptor emission intensity at 618 nm by the donor emission intensity at 460 nm. Values were corrected by subtracting the mean BRET ratio measured in the corresponding no-ligand control wells and are reported in milliBRET units (mBU), calculated by multiplying the background-corrected BRET ratio by 1,000. NanoBRET assay vectors were a gift from Promega Corporation (Addgene plasmids #238827, #237019, #238570).

Statistical significance of the changes in BRET ratios displayed in **Figure 2 and Table 1** was determined using a two-sided unpaired Welch’s t-test in SciPy [42] to account for unequal sample variances. For all statistical analyses, compound treatments were compared directly against the corresponding MG-132 alone baseline control. The raw data can be found in **Supplementary Table 3.**

### Mechanistic validation of VHL and CRL-mediated target modulation

Whole-cell lysates were prepared from peptide-transfected and/or siRNA-transfected A549 or HCT116 cells, as appropriate for KRAS- or BRD4-related experiments. Cells were lysed in NP-40 lysis buffer (50 mM Tris-HCl pH 8.0, 150 mM NaCl, 1% NP-40) supplemented with protease inhibitors (Complete Mini, Roche) and phosphatase inhibitors (PhosSTOP, Roche). Proteins were resolved by SDS-PAGE on 4-12% Bis-Tris gels and transferred to PVDF membranes. Membranes were cut into strips corresponding to the expected molecular weights of the proteins of interest and probed with the following primary antibodies: BRD4 (Bethyl Laboratories, RRID:AB_2631450), K-RAS (Abcam, RRID:AB_2884935), VHL (Santa Cruz Biotechnology, RRID:AB_2215955), MYC (Abcam, RRID:AB_731658) and GAPDH (Santa Cruz Biotechnology, RRID:AB_10167668). Horseradish peroxidase-conjugated, isotype-specific secondary antibodies were used for chemiluminescent detection. For proteins of similar molecular weights, membrane strips were treated with a stripping buffer (Thermo Fisher Scientific, 46430) and sequentially reprobed with the relevant antibodies. The results from these analyses can be found in **Figure 3** and **Table 3**. The full blots can be found in **Supplementary Figures 3 and 4** and are representative of independent biological triplicates.

### Neuroblastoma tumoroid culture and BRD4 degradation assays

The MYCN-amplified patient-derived xenograft (PDX)-derived neuroblastoma tumoroid models LU-NB-2 and LU-NB-3 have previously been described and retain the molecular and tumorigenic characteristics of the originating patient tumours [43]. All experiments were performed under the relevant ethical approvals.

Tumoroids were maintained in Neuroblastoma tumoroid Medium (NBOM) consisting of DMEM (low glucose, GlutaMAX™ Supplement, pyruvate; Life Technologies, Cat. 21885108), Ham’s F-12 Nutrient Mix (GlutaMAX™; Life Technologies, Cat. 31765027), B-27™ Supplement (50×, minus vitamin A; Life Technologies, Cat. 12587010), N-2 Supplement (100×; Life Technologies, Cat. 17502048), Penicillin-Streptomycin (10,000 U/mL; Life Technologies, Cat. 15140122), recombinant human EGF (PeproTech, Cat. AF-100-15), FGF-basic (PeproTech, Cat. 100-18B), IGF-I (PeproTech, Cat. 100-11), PDGF-AA (PeproTech, Cat. 100-13A), and PDGF-BB (PeproTech, Cat. 100-14B).

For degradation assays, tumoroids were dissociated into single-cell suspensions and seeded in ultra-low attachment 12-well plates at 3 × 10⁵ cells per well in 1 mL NBOM. Cells were allowed to reform tumoroids for 48 h before treatment. Where indicated, VHL knockdown was performed by siRNA transfection 24 h before peptide treatment. The BRD4-targeting macrocycles B195 and B938 were delivered using SAINT-Protein transfection reagent (Synvolux Therapeutics). Peptide-SAINT complexes were prepared in PBS according to the manufacturer’s protocol, incubated for 5-10 min at room temperature and added directly to the cultures to achieve a final peptide concentration of 10 µM. Control wells received the corresponding amount of SAINT reagent alone. Cells were incubated with the peptide complexes for 24 h. Where indicated, the neddylation inhibitor MLN4924 (Selleckchem, S7109) was added 4 h before sample collection, and the BRD4 degrader MZ-1 was included as a positive control.

Following treatment, tumoroids were collected by centrifugation, washed once with PBS and lysed in RIPA buffer supplemented with protease and phosphatase inhibitors. Protein concentrations were determined using the Pierce™ BCA Protein Assay Kit (Thermo Fisher Scientific), and equal amounts of protein were separated on 3-8% NuPAGE™ Tris-Acetate gels (Thermo Fisher Scientific) using NuPAGE™ Tris-Acetate SDS running buffer. Proteins were transferred onto 0.45 µm nitrocellulose membranes using the Power Blotter XL system (Thermo Fisher Scientific) with the High Molecular Weight transfer program. Membranes were blocked in Intercept® TBS Blocking Buffer (LI-COR Biosciences) and incubated overnight at 4°C with primary antibodies against BRD4 (Bethyl Laboratories, A301-985A100), N-Myc (Cell Signaling Technology, 51705), and β-actin (Cell Signalling Technology, 4970). Following washing with TBST, membranes were incubated with IRDye secondary antibodies (LI-COR Biosciences) for 1 h at room temperature. Fluorescent signals were acquired using an Odyssey imaging system (LI-COR Biosciences), and band intensities were quantified using Image Studio software. BRD4 and N-Myc signal intensities were normalised to β-actin.

The full blots can be found in **Supplementary Figures 5-8**.

## Data availability

All data are available at: https://zenodo.org/uploads/14065843

## Code availability

EvoBind-multimer and an easy-to-use Colab notebook are available here: https://github.com/patrickbryant1/EvoBind-multimer

## Funding

This study was supported by the SciLifeLab & Wallenberg Data Driven Life Science Program (KAW 2020.0239, P.B). This work was supported by the Swedish Cancer Society (23 2708 Pj, K.K.). A.B.’s salary is supported by Worldwide Cancer Research (grant reference 26-0033). Work in the OS laboratory was further supported by the Swedish Cancer Society, the Swedish Childhood Cancer Fund, the Swedish Research Council, Karolinska Institutet and Radiumhemmets Research Foundation. The computing power was enabled by the Berzelius resource provided by the Knut and Alice Wallenberg Foundation at the National Supercomputer Centre with project IDs Berzelius-2023-267, Berzelius-2024-78, Berzelius-2024-292, Berzelius-2025-41 and Berzelius-2025-247 (P.B).

## Contributions

P.B. conceived and designed the study, developed EvoBind-multimer, generated and selected binder designs, prepared the figures, and wrote the initial manuscript draft. A.B. and O.S. performed the live-cell NanoBRET and mechanistic analyses. K.W. and K.K. performed the tumoroid experiments, with support from D.B. D.D. coordinated peptide logistics, developed the GitHub repository and Google Colab notebooks for distribution, and analysed structural binding modes in coordination with P.B. All authors reviewed and edited the manuscript.

## Conflicts of interest

P.B. is a shareholder in Cyclic Therapeutics, a company that designs and develops therapeutic peptides.

## Supplementary information

### Supplementary Figures

**Supplementary Figure 1.**
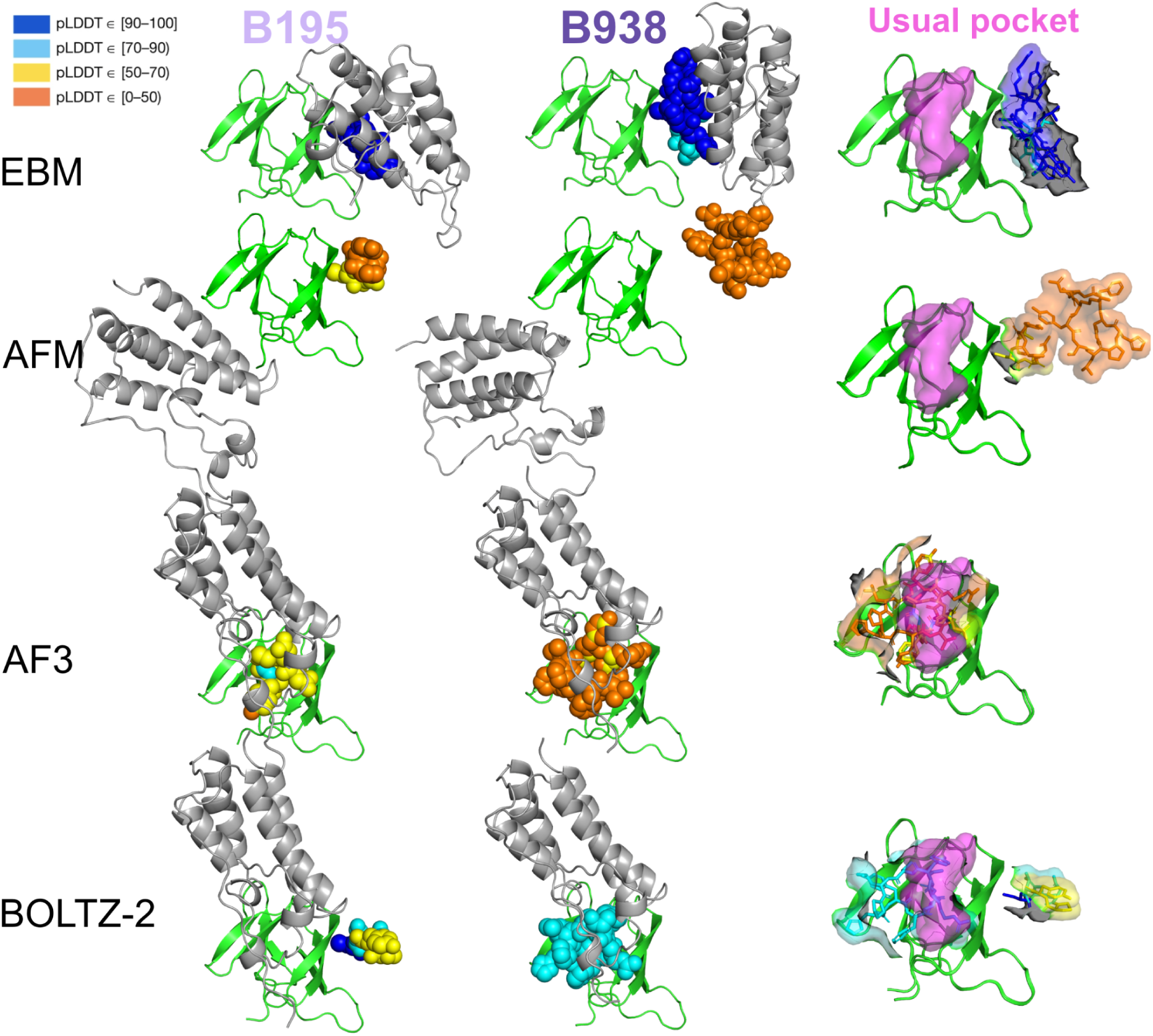
Cross-validation of the top 2 ternary complexes of BRD4 with different AI structure prediction methods. The two target proteins, VHL (green) and BRD4 (grey), are shown in cartoon format. The cyclic peptide molecular glue is shown as spheres colored by pLDDT (see legend). All structures are superimposed on VHL for easier comparison. All adversarial predictions fail to locate the molecular glue and/or BRD4, showing a pattern of overfitting towards the front side of VHL, which is the only binding site of all known small-molecule molecular glues. In the third column, the usual binding pocket surface of known small-molecule glues (from PDB ID 8QU8) is shown in magenta, while the two peptide glues from each model are colored by pLDDT and shown together as sticks and surfaces.

**Supplementary Figure 2.**
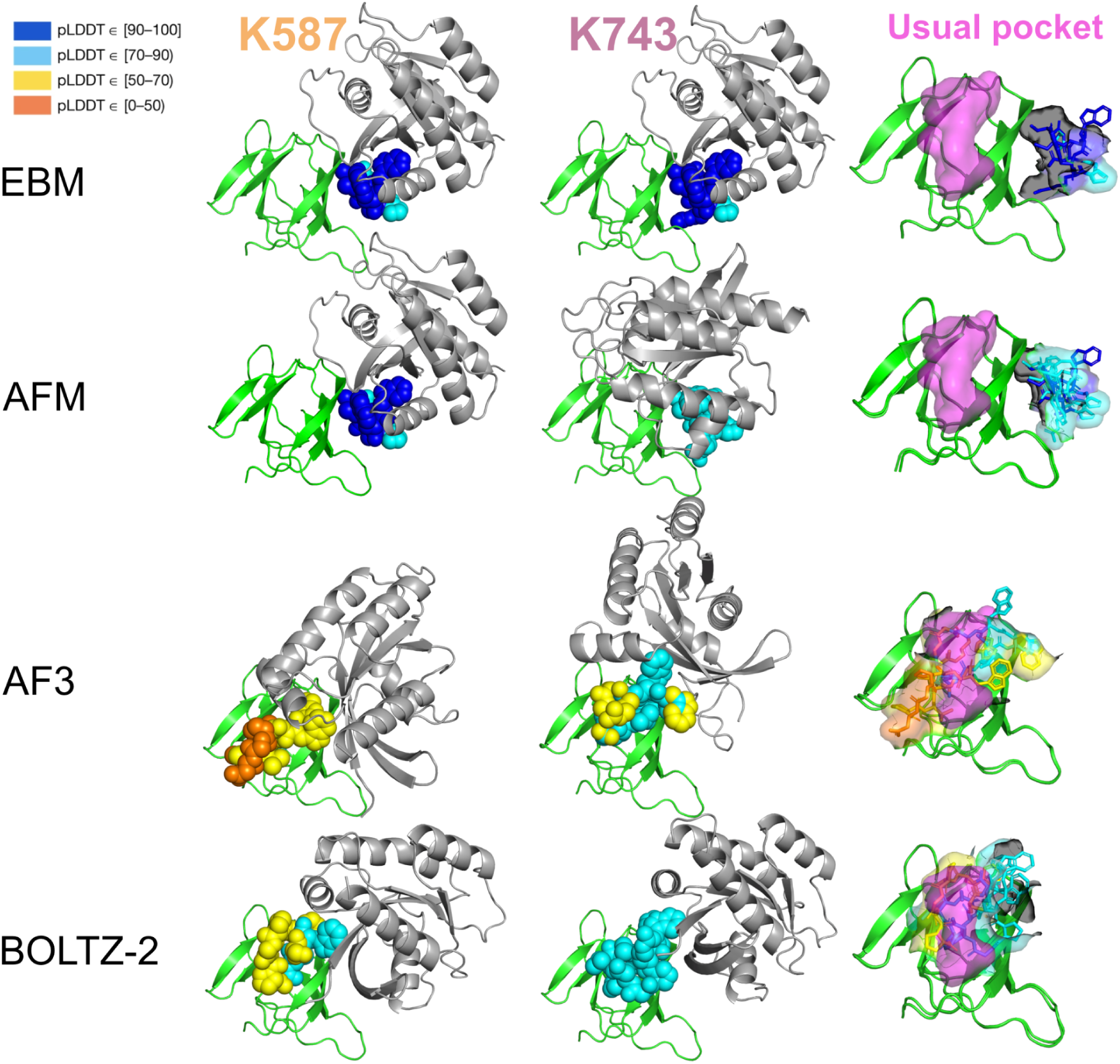
Cross-validation of the top 2 ternary complexes of KRAS with different AI structure prediction methods. The two target proteins, VHL (green) and KRAS (grey), are shown as cartoons. The cyclic peptide molecular glue is shown as spheres colored by pLDDT (see legend). All structures are superimposed on VHL for easier comparison. All adversarial predictions fail to locate the molecular glue and/or KRAS, showing a pattern of overfitting towards the front side of VHL, which is the only binding site of all known small-molecule molecular glues. In the third column, the usual binding pocket surface of known small-molecule glues (from PDB ID 8QU8) is shown in magenta, while the two peptide glues from each model are colored by pLDDT and shown together as sticks and surfaces.

**Supplementary Figure 3.**
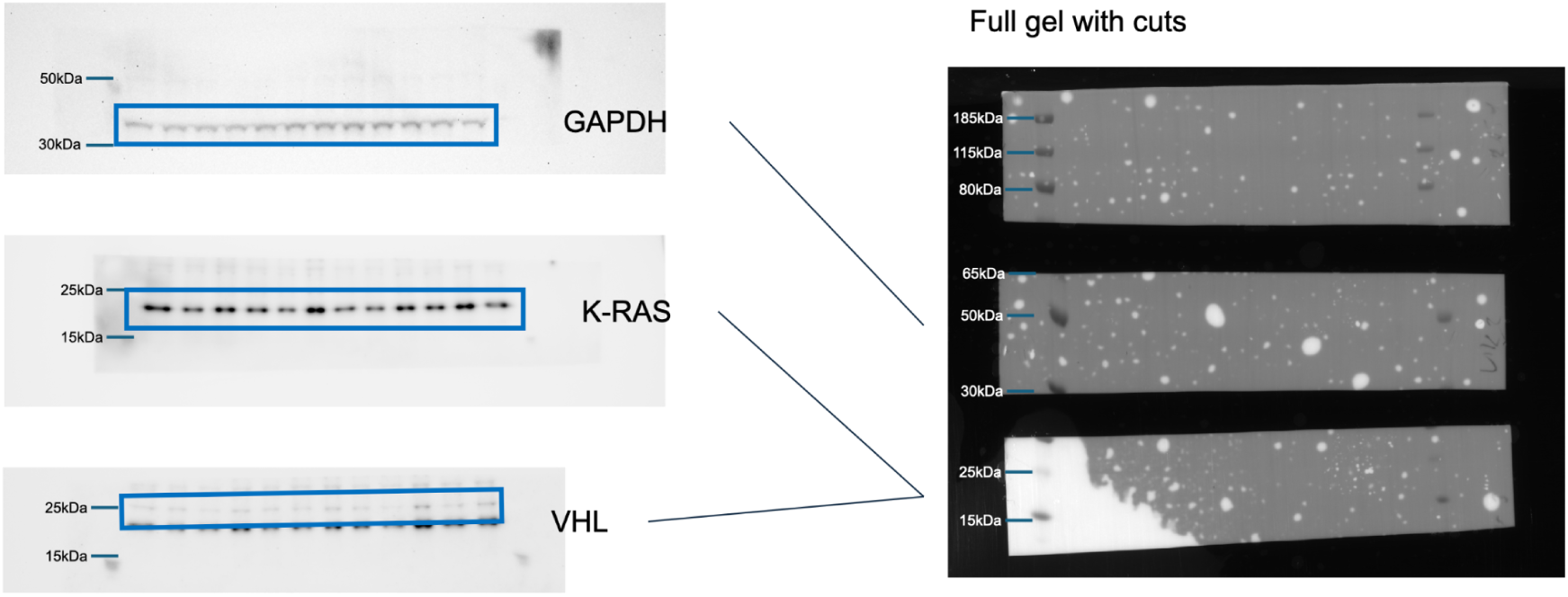
Full uncropped membrane blot images for KRAS mechanistic validation (refers to Figure 3a). Uncropped chemiluminescence images (left) and corresponding physical membrane cuts (right) for the immunoblot analysis shown in **Figure 3a**. To optimise antibody incubation across distinct molecular weight ranges, PVDF membranes were cut horizontally into three sections before probing (right panel). The middle membrane section (∼30–65 kDa) was probed for the GAPDH loading control (∼36 kDa). Because KRAS (∼21 kDa) and VHL (∼18-24 kDa) migrate within the same lower molecular weight region (<30 kDa), the bottom membrane strip was sequentially probed and reprobed, analysing the target with the weaker signal first before subsequent antibody re-incubation. Blue boxes indicate the cropped membrane areas presented in **Figure 3a**. Molecular weight ladder positions (kDa) are indicated on the left of each panel. The second VHL band is likely the probable p19 isoform or a non-specific band.

**Supplementary Figure 4.**
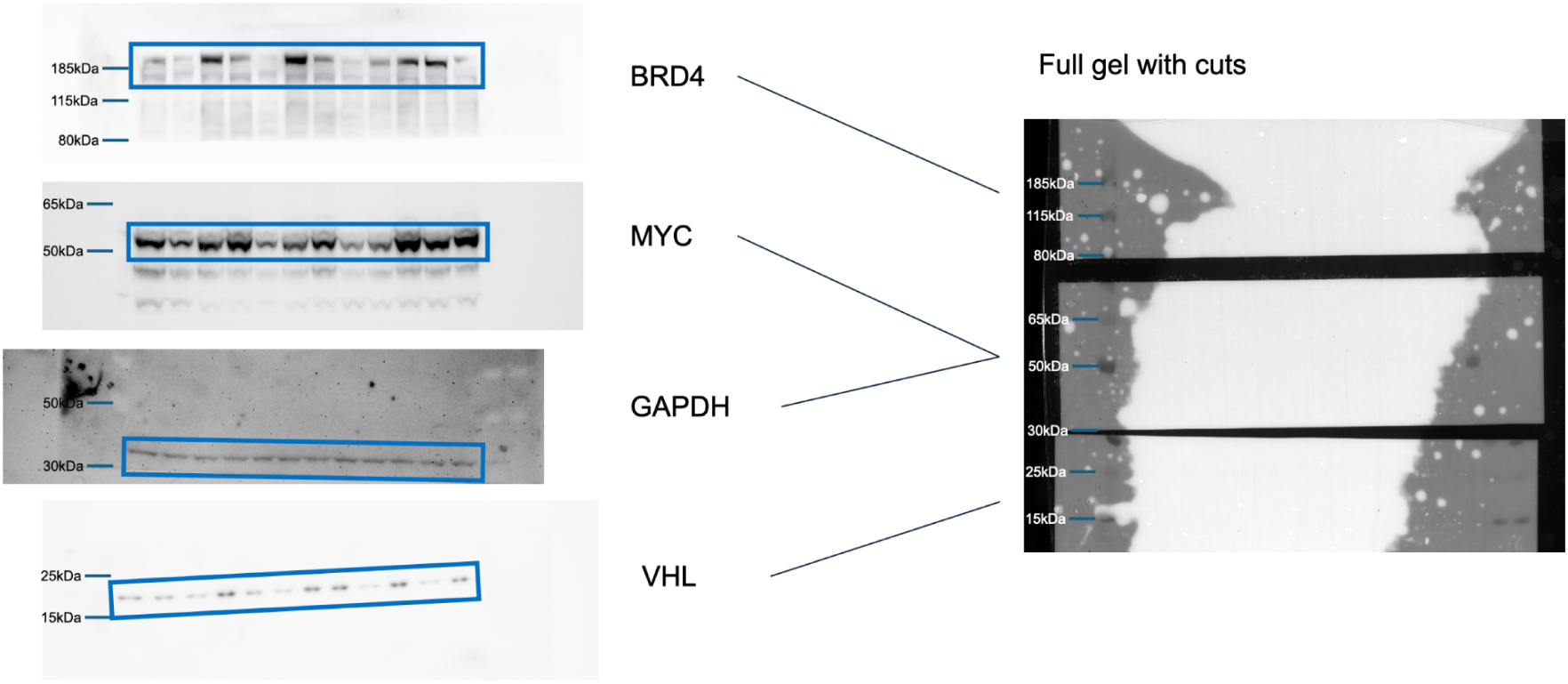
Full uncropped membrane blot images for BRD4 mechanistic validation (refers to Figure 3b). Uncropped chemiluminescence images (left) and corresponding physical membrane cuts (right) for the immunoblot analysis shown in **Figure 3b**. To optimise antibody incubation across distinct molecular weight ranges, PVDF membranes were cut horizontally into three sections before probing (right panel). The top membrane section (>80 kDa) was probed for BRD4. The middle membrane section (∼30–80 kDa) encompasses the regions probed for downstream MYC and the GAPDH loading control. The bottom membrane section (<30 kDa) was probed for VHL. Blue boxes indicate the explicitly cropped membrane areas presented in Figure 3b. Molecular weight ladder positions (kDa) are indicated on the left of each panel.

**Supplementary Figure 5.**
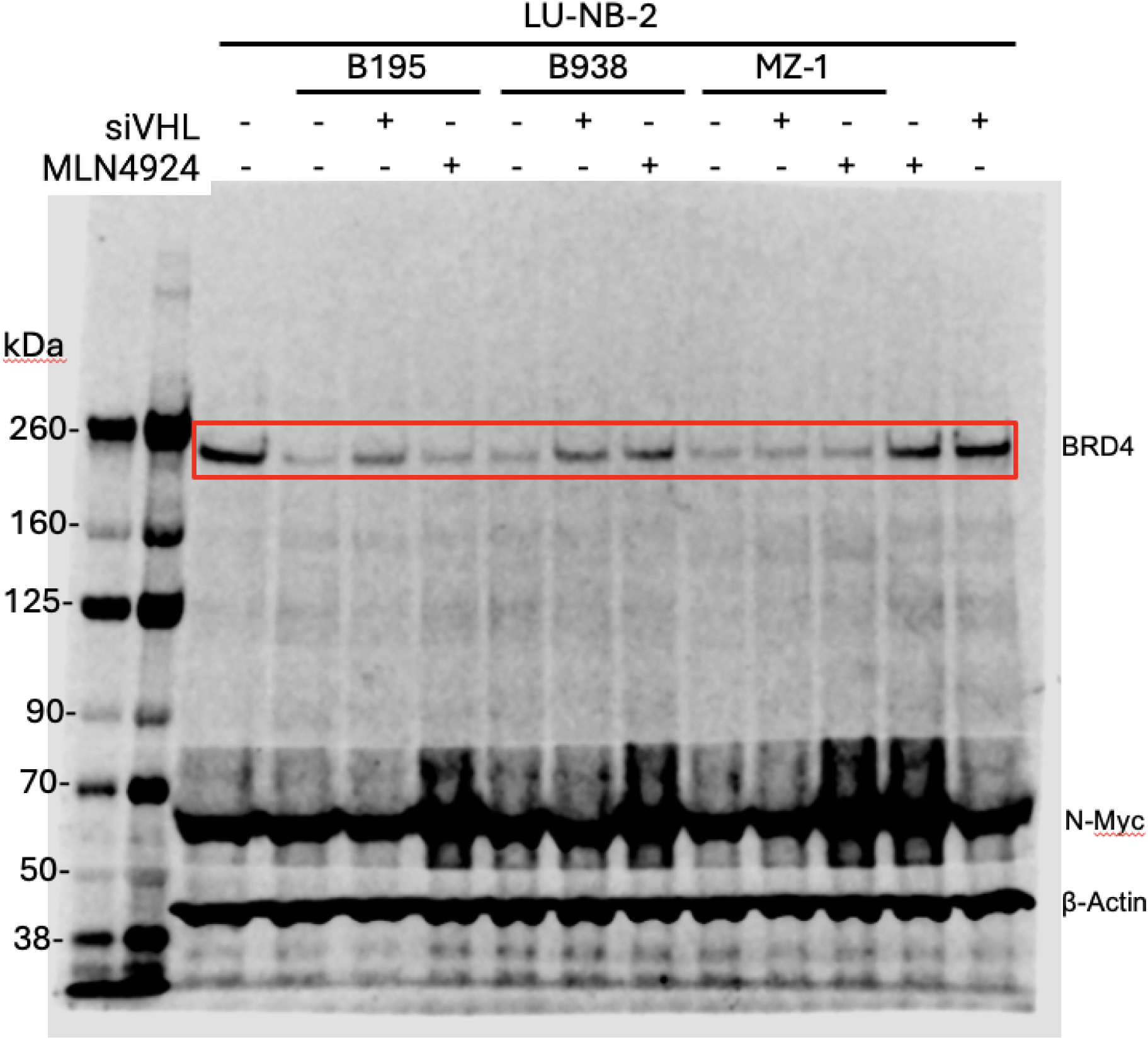
Full uncropped membrane blot image for BRD4 degradation in LU-NB-2 tumoroids. Uncropped chemiluminescence membrane image for the immunoblot analysis performed in patient-derived xenograft LU-NB-2 neuroblastoma tumoroids (shown cropped from the region marked in red in **Figure 4b**). Tumoroids were treated with degraders (B195, B938) or reference PROTAC (MZ-1) in the presence or absence of VHL knockdown (siVHL) or neddylation inhibition (MLN4924) to confirm E3 ligase and proteasome pathway dependency. Immunoblotting was performed for BRD4 (∼200 kDa), downstream target N-Myc (∼60–65 kDa), and β-Actin (∼42 kDa) as a loading control. The red box highlights the explicitly cropped BRD4 region presented in **Figure 4b**. Protein molecular weight marker positions (kDa) are indicated on the left side of the gel.

**Supplementary Figure 6.**
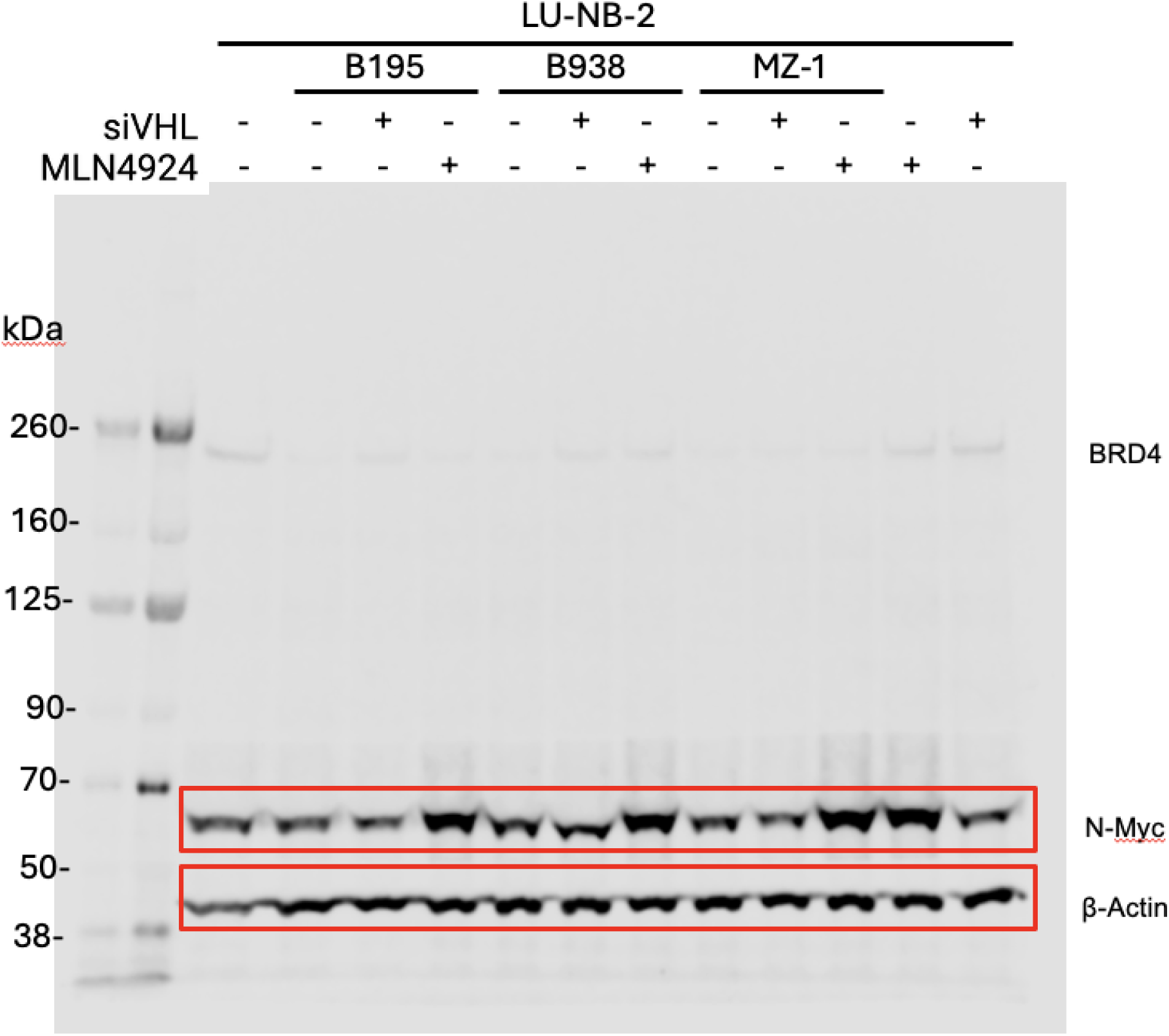
Full uncropped membrane blot images for downstream N-Myc and loading control β-Actin in LU-NB-2 tumoroids. Uncropped chemiluminescence membrane image for the immunoblot analysis performed in patient-derived xenograft LU-NB-2 neuroblastoma tumoroids (shown cropped from the region marked in red in **Figure 4b**). Tumoroids were treated with degraders (B195, B938) or reference PROTAC (MZ-1) in the presence or absence of VHL knockdown (siVHL) or neddylation inhibition (MLN4924). Immunoblotting was performed for the downstream oncogenic driver N-Myc (∼60-65 kDa) and β-Actin (∼42 kDa) as an internal loading control (faint BRD4 signal at ∼200 kDa is also visible in the upper region of the membrane). Red boxes highlight the explicitly cropped membrane regions presented in **Figure 4b**. Protein molecular weight marker positions (kDa) are indicated on the left side of the gel.

**Supplementary Figure 7.**
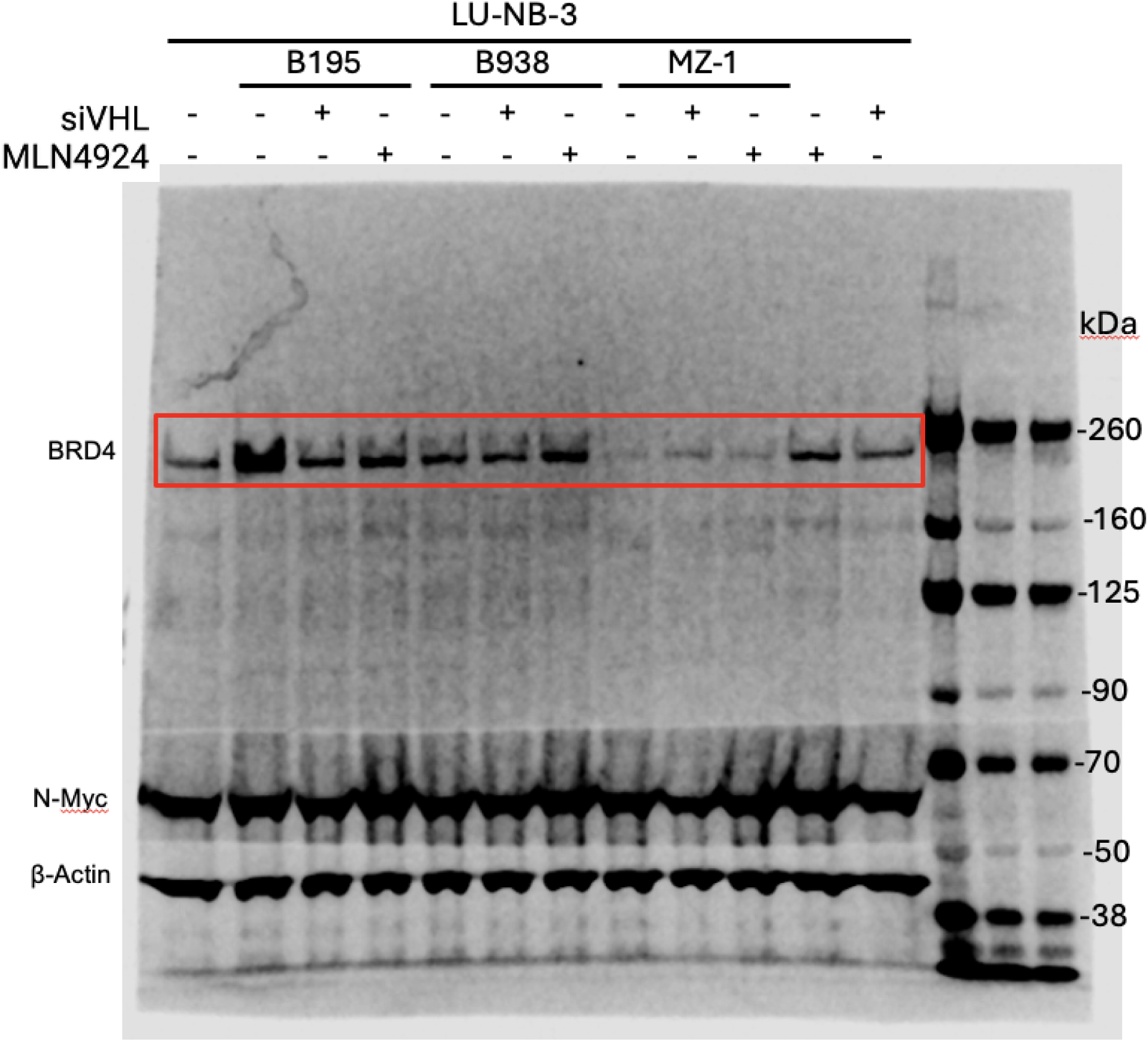
Full uncropped membrane blot image for BRD4 degradation in LU-NB-3 tumoroids. Uncropped chemiluminescence membrane image for the immunoblot analysis performed in patient-derived xenograft LU-NB-2 neuroblastoma tumoroids (shown cropped from the region marked in red in **Figure 4b**). Tumoroids were treated with degraders (B195, B938) or reference PROTAC (MZ-1) in the presence or absence of VHL knockdown (siVHL) or neddylation inhibition (MLN4924) to confirm E3 ligase and proteasome pathway dependency. Immunoblotting was performed for BRD4 (∼200 kDa), downstream target N-Myc (∼60-65 kDa), and β-Actin (∼42 kDa) as a loading control. The red box highlights the explicitly cropped BRD4 region presented in **Figure 4b**. Protein molecular weight marker positions (kDa) are indicated on the right side of the gel.

**Supplementary Figure 8.**
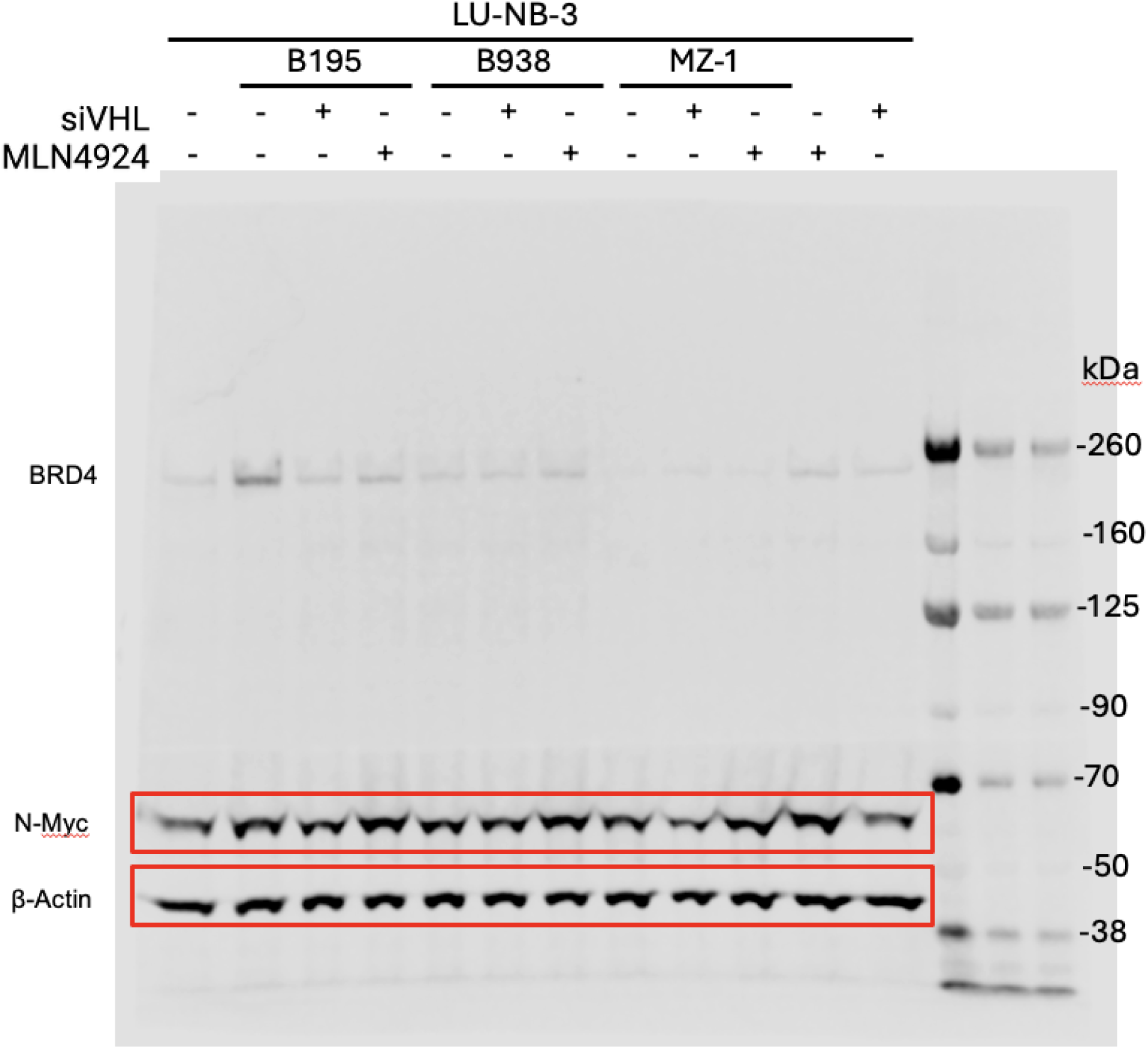
Full uncropped membrane blot images for downstream N-Myc and loading control β-Actin in LU-NB-2 tumoroids. Uncropped chemiluminescence membrane image for the immunoblot analysis performed in patient-derived xenograft LU-NB-2 neuroblastoma tumoroids (shown cropped from the region marked in red in **Figure 4b**). Tumoroids were treated with degraders (B195, B938) or reference PROTAC (MZ-1) in the presence or absence of VHL knockdown (siVHL) or neddylation inhibition (MLN4924). Immunoblotting was performed for the downstream oncogenic driver N-Myc (∼60-65 kDa) and β-Actin (∼42 kDa) as an internal loading control (faint BRD4 signal at ∼200 kDa is also visible in the upper region of the membrane). Red boxes highlight the explicitly cropped membrane regions presented in **Figure 4b**. Protein molecular weight marker positions (kDa) are indicated on the right side of the gel.

### Supplementary Tables

**Supplementary Table 1.**
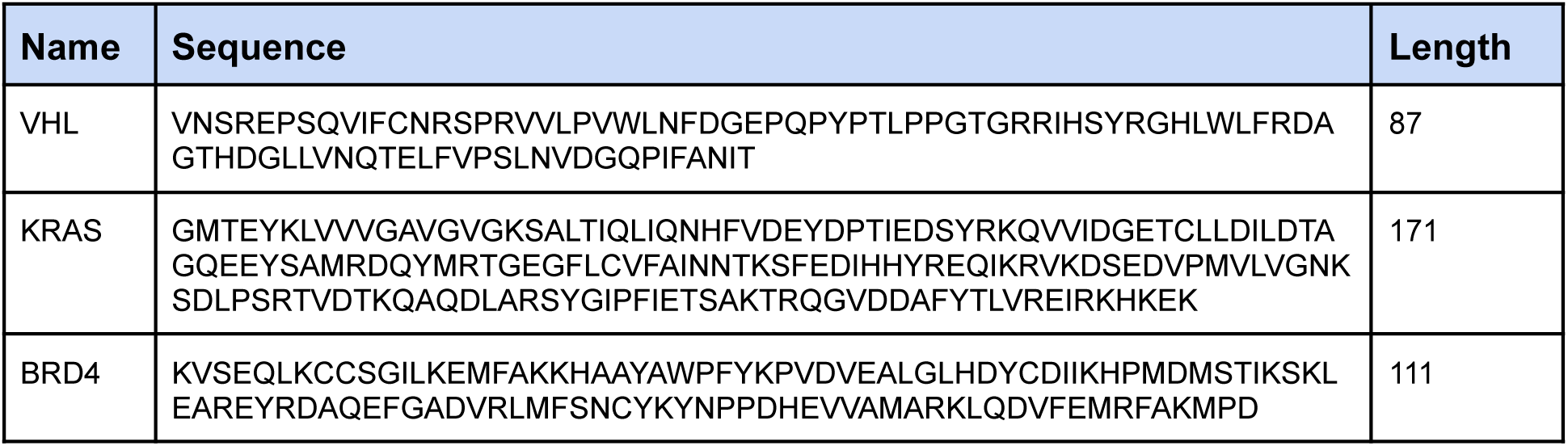
Target sequences and lengths.

**Supplementary Table 2.**
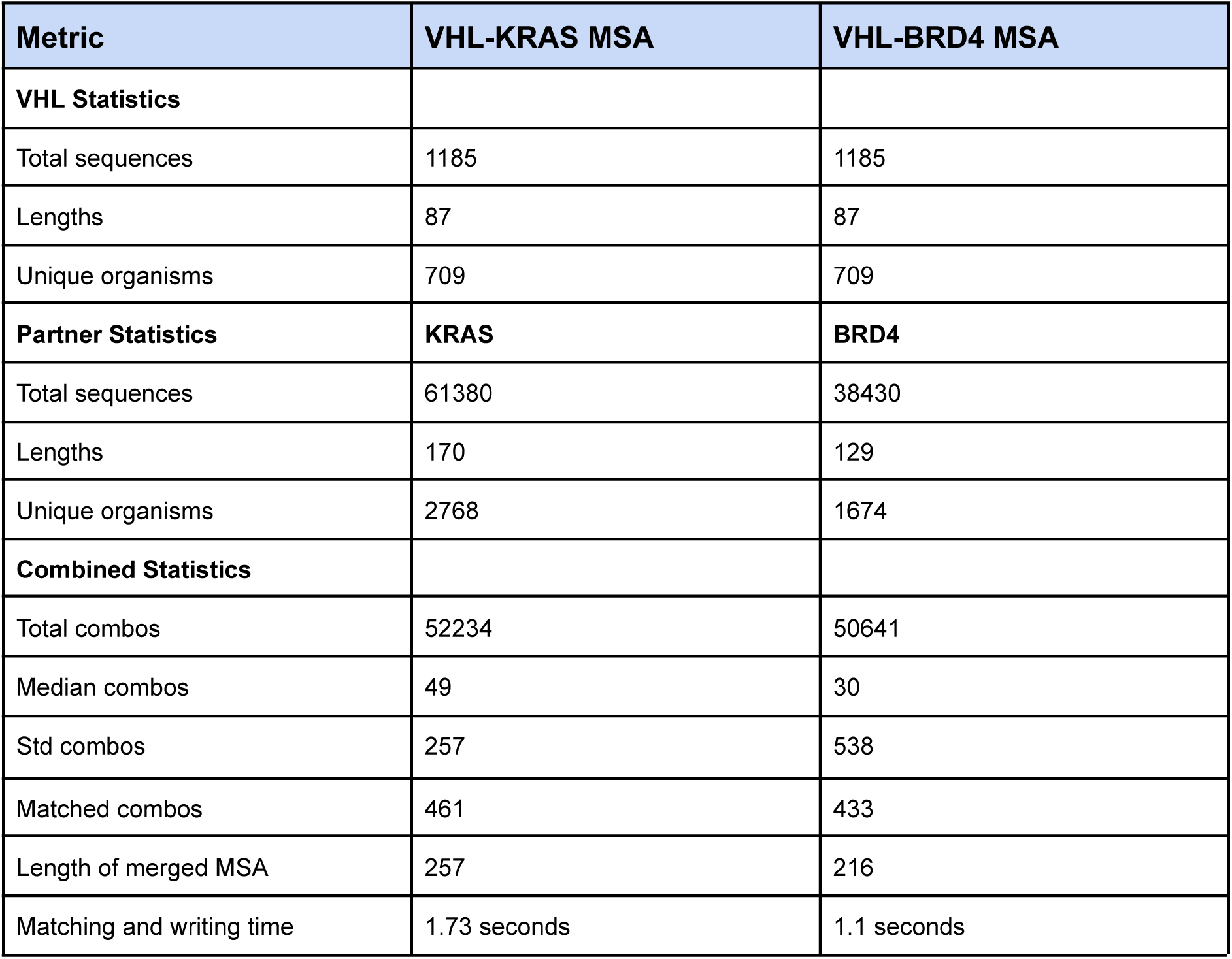
MSA statistics. Comparison of VHL-KRAS and VHL-BRD4 MSAs.

**Supplementary Table 3.** Live-cell NanoBRET results. The target (KRAS or BRD4) indicates the target protein that is not VHL; the condition indicates the treatment condition, and R1-R3 the result from three independent biological replicates.

| Target | Condition | R1 | R2 | R3 |
| --- | --- | --- | --- | --- |
| KRAS | DMSO | 6.967854 | 6.572644 | 6.468197 |
| KRAS | + MG-132 | 7.724976 | 6.780998 | 5.98654 |
| KRAS | K587 | 7.679598 | 7.814742 | 7.401347 |
| KRAS | K587 + MG-132 | 9.212484 | 8.537298 | 10.31596 |
| KRAS | K743 | 7.312243 | 8.270388 | 8.022133 |
| KRAS | K743 + MG-132 | 9.128841 | 8.924931 | 9.073969 |
| KRAS | LC-2 | 7.623117 | 7.549035 | 7.467741 |
| KRAS | LC-2 + MG-132 | 11.60486 | 10.31186 | 9.002152 |
| BRD4 | DMSO | 6.571624 | 6.416809 | 6.454133 |
| BRD4 | + MG-132 | 7.483153 | 7.186465 | 7.452235 |
| BRD4 | B195 | 7.096591 | 6.888152 | 7.461119 |
| BRD4 | B195 + MG-132 | 9.187312 | 8.832788 | 8.419358 |
| BRD4 | B938 | 7.197575 | 7.271434 | 7.674745 |
| BRD4 | B938 + MG-132 | 8.89198 | 9.156976 | 9.233934 |
| BRD4 | MZ1 | 7.478143 | 7.10029 | 7.353402 |
| BRD4 | MZ1 + MG-132 | 9.310341 | 8.776573 | 8.396436 |

## Notes

https://zenodo.org/uploads/14065843

